# DNA Calorimetric Force Spectroscopy at Single Base Pair Resolution

**DOI:** 10.1101/2024.04.29.591589

**Authors:** P. Rissone, M. Rico-Pasto, S. B. Smith, F. Ritort

**Author notes:** Contributing authors.

## Abstract

DNA hybridization is a fundamental reaction with wide-ranging applications in biotechnology. The nearest-neighbor (NN) model provides the most reliable description of the energetics of duplex formation. Most DNA thermodynamics studies have been done in melting experiments in bulk, of limited resolution due to ensemble averaging. In contrast, single-molecule methods have reached the maturity to derive DNA thermodynamics with unprecedented accuracy. We combine single-DNA mechanical unzipping experiments using a temperature jump optical trap with machine learning methods and derive the temperature-dependent DNA energy parameters of the NN model. In particular, we measure the previously unknown ten heat-capacity change parameters **Δ*C***_***p***_, relevant for thermodynamical predictions throughout the DNA stability range. Calorimetric force spectroscopy establishes a groundbreaking methodology to accurately study nucleic acids, from chemically modified DNA to RNA and DNA/RNA hybrid structures.

---

Nucleic acid (NA) thermodynamics is essential to understand duplex formation, where two single strands of DNA or RNA hybridize to form a double helix. Hybridization is a crucial process for genome maintenance with many biotechnological applications, from PCR amplification to gene editing ^1^ and DNA origami ^2,3^. Accurate knowledge of the thermodynamic energy parameters of NA hybridization is necessary for developing better protocols, often involving heating and cooling cycles for dissociating and hybridizing complementary strands. Most hybridization studies involve calorimetric melting experiments in bulk, where signals such as heat, UV light absorbance, and fluorescence are measured over samples typically containing 10^10^ molecules in aqueous solutions ^4,5^.

Despite recent progress, basic questions about NA hybridization remain unanswered, such as the nature of the transition state and duplex stability far from standard conditions. Examples are extreme cold and high temperatures, molecular condensates, and confined spaces. Thermodynamic predictions far from standard conditions require far-fetched extrapolations of the currently known energy parameters. The heat-capacity change at constant pressure, Δ*C*_*p*_, is crucial for NA formation. Δ*C*_*p*_ quantifies the temperature dependence of enthalpy (Δ*H*) and entropy (Δ*S*) contributions to the free energy of hybridization through the thermodynamic relations at constant pressure, Δ*C*_*p*_ = *∂*Δ*H/∂T* = *T∂*Δ*S/∂T* . From a microscopic viewpoint, Δ*C*_*p*_ relates to the change in the number of degrees of freedom, Δ*n*, in hybridization reactions according to the equipartition law, Δ*C*_*p*_ = *k*_*B*_Δ*n/*2, with Δ*n* = 1 per cal mol^*−*1^K^*−*1^ unit of Δ*C*_*p*_. It has been suggested that the most significant contribution to Δ*C*_*p*_ in duplex formation occurs in the alignment of the complementary single-strands upon hybridization ^6^.

The temperature dependence of the enthalpy and entropy of DNA hybridization has been neglected for a long time. The assumption Δ*C*_*p*_ = 0 was mainly adopted during the first scanning calorimetry studies that could not detect Δ*C*_*p*_ ^7^. Over time, improvements in calorimetric measurements ^8^ pointed out the significant role Δ*C*_*p*_ played in DNA hybridization. During the last decades, bulk ^9–15^ and single-molecule ^16–18^ experiments assessed the effects of temperature, yielding Δ*C*_*p*_ values per bp spanning two orders of magnitude depending on the experimental condition, the technique used and the DNA sequence. Single-molecule methods such as atomic force microscopy ^19,20^, FRET ^21^, and optical tweezers ^22,23^ have now reached the maturity to address such challenges. While force spectroscopy derives free energy differences (Δ*G*) from work measurements, single-molecule FRET does it from the lifetimes of single NA hairpins permits measuring the folding free energy landscape along the reaction coordinate defined by the number of released nucleotides during the unzipping process ^25,26^. The unzipping reaction finds applications in the footprinting of DNA-binding restriction enzymes ^27^, transcription factors ^28,29^, and peptides ^30^. Mechanical unzipping has also permitted the design of calipers for measuring molecular distances ^31^. Here, we derive the elusive heat capacity change Δ*C*_*p*_ for the different DNA nearest-neighbor base pair motifs.

DNA’s thermodynamic stability (Δ*G*) results from the compensation of the favorable Δ*H* and the unfavorable Δ*S* of folding, Δ*G* = Δ*H − T* Δ*S*, with *T* the temperature. For Δ*C*_*p*_ = 0, Δ*H* and Δ*S* are *T* -independent, and Δ*G* is linear with *T* . For Δ*C*_*p*_*≠* 0, enthalpy-entropy compensation makes Δ*H* and *T* Δ*S* of comparable magnitude masking deviations of Δ*G* from a *T* -linear behavior. Potentially, one could derive the temperature-dependent Δ*S* from Δ*G* using the relation Δ*S* = *−∂*Δ*G/∂T* . However, this method is imprecise due to strong compensation between Δ*H* and *T* Δ*S*. To determine Δ*C*_*p*_, it is convenient to measure either enthalpy or entropy contributions independently of Δ*G*. We introduce a method to accurately derive DNA thermodynamics by directly measuring the temperature-dependent entropy of hybridization Δ*S* using calorimetric force spectroscopy with optical tweezers ^17^. We apply the Clausius-Clapeyron equation to single-molecule experiments ^32^ and combine it with a tailored machine-learning algorithm. This approach allows us to derive the entropies, enthalpies, and Δ*C*_*p*_ parameters of hybridization at single base pair resolution in the nearest-neighbor (NN) model ^33,34^.

According to the NN model, the duplex’s free energy, entropy, and enthalpy equal the sum of all contributions of the adjacent nearest-neighbor base pairs (NNBP) along the sequence. The four distinct canonical Watson-Crick base pairs generate sixteen possible NNBP combinations (e.g., AG/TC, meaning that 5^*′*^*−*AG*−*3^*′*^ hybridizes with 3^*′*^*−*TC*−*5^*′*^) with their corresponding energy parameters. The sixteen parameters reduce to ten due to strand complementarity symmetry (e.g., AG/TC equals CT/GA), further reducing to eight from circular symmetry relations ^35–37^. The ten NNBP DNA parameters have been derived at 37^*°*^C from melting experiments of short DNA duplexes by many laboratories worldwide ^38–42^ and unified by Santalucia *et al*. ^43^ in the so-called *Unified Oligonucleotide* (UO) set. In the last decade, the NNBP free energy parameters have been derived from reversible work measurements in mechanical unzipping experiments of DNA and RNA hairpins at room temperature (298K) ^26,37,44^. These energy parameters are used by most secondary structure prediction tools, such as Mfold ^45^, Vienna package ^46^, uMelt ^47^, among others. Here, we apply calorimetric force spectroscopy to measure the NNBP energy parameters in the temperature range 7 *−* 40^*°*^C and derive the Δ*C*_*p*_ values.

## Results

We used a temperature-jump optical trap (Fig. 1A and Sec. 1, Methods) to unzip a 3593bp (*≈* 3.6kbp) DNA hairpin made ending in a GAAA tetraloop. Pulling experiments were conducted at temperatures 7 *−* 42^*°*^C at 1M NaCl in 10mM Tris-HCl buffer (pH 7.5). Figure 1B shows the measured force-distance curves (FDCs). They exhibit a sawtooth pattern upon increasing the trap-pipette distance, *λ*, until the hairpin unzips completely and the elastic response of the released single-stranded DNA (ssDNA) is measured (rightmost part of Fig. 2B). Upon increasing *T*, the hairpin unzips at progressively lower forces (horizontal dashed lines in Fig.1B) from *∼* 20pN at 7^*°*^C to *∼* 14pN at 42^*°*^C. This indicates that the DNA stability decreases with *T*, and the energy parameters of the NN model are temperature dependent: the higher the temperature, the lower the hairpin’s free energy of hybridization. Moreover, the molecular extension of the ssDNA at a given force increases with *T*, yielding a total of *∼* +500nm between 7^*°*^ and 42^*°*^C (Fig. 2B, horizontal grey double arrow).

**Fig. 1.**
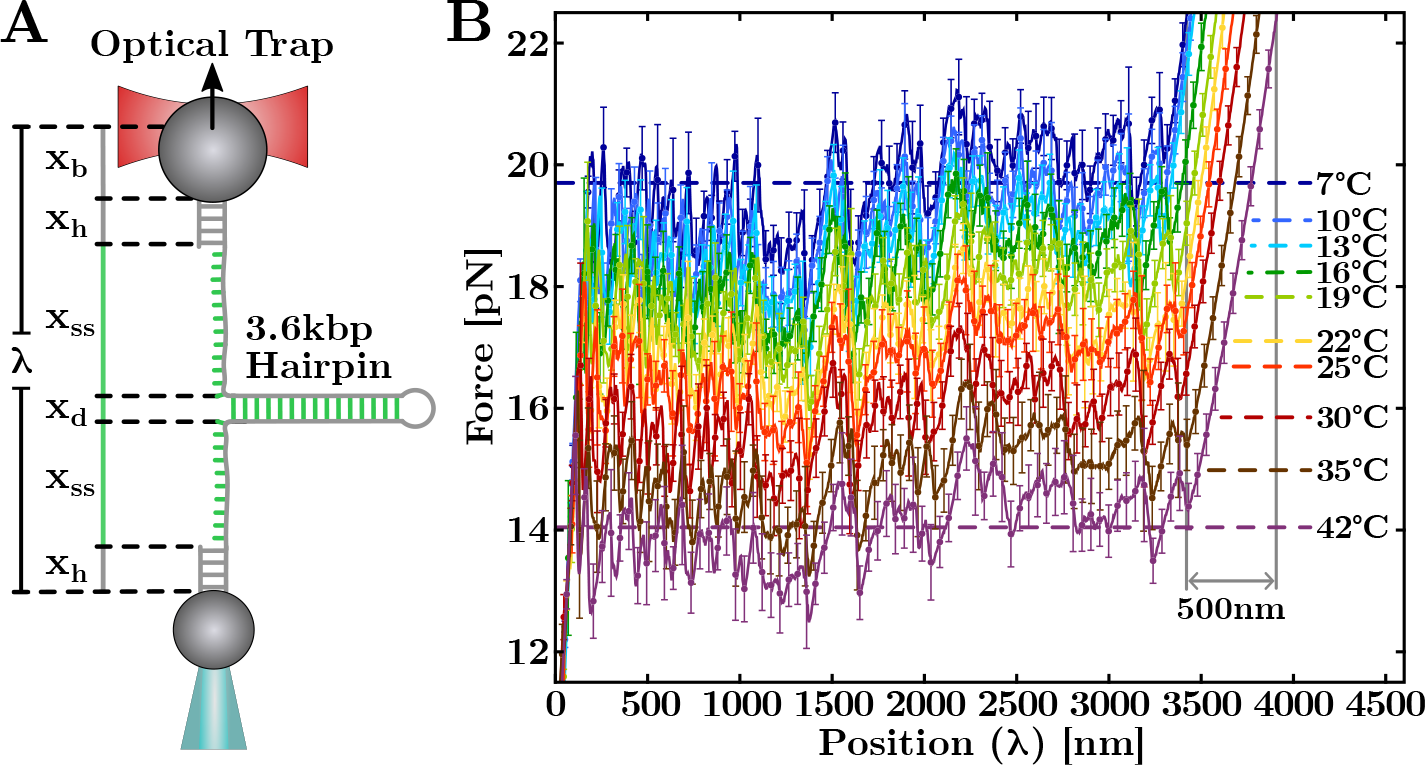
Single-molecule calorimetric force-spectroscopy. (**A**) Experimental setup (Sec. 1, Methods) showing each component of the (measured) total trap-pipette distance, *λ*. (**B**) Temperature dependence of FDCs obtained by pulling the 3.6kbp DNA hairpin. At each *T*, the FDC results from averaging over 5 *−* 6 molecules for 40 *−* 50 unzipping/rezipping cycles. The error bars (plotted for a fraction of the total data points) show the molecule-to-molecule variability.

**Fig. 2.**
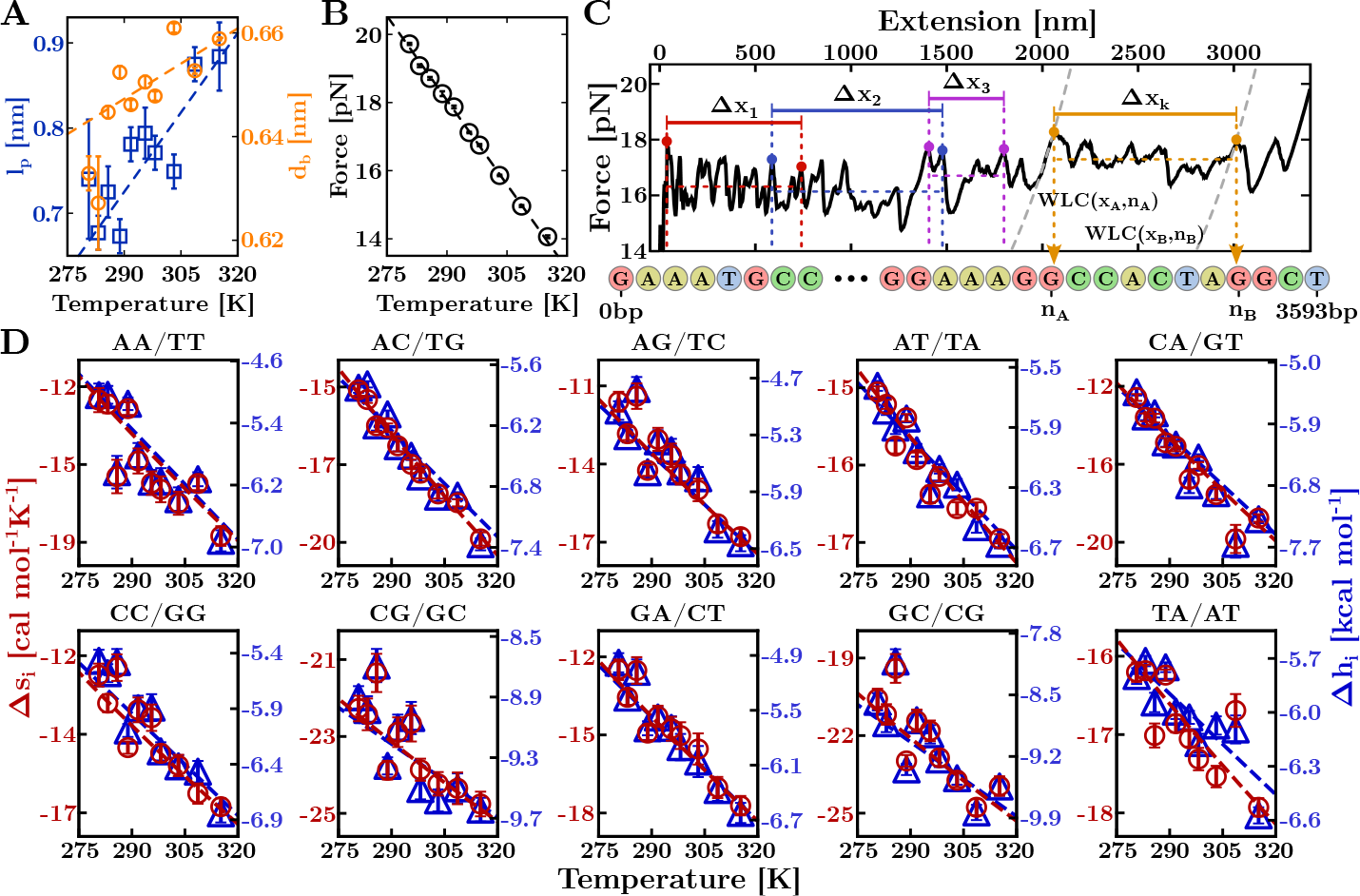
NNBP entropies and enthalpies. (**A, B**) *T* -dependence of the persistence length, *l*_*p*_, (blue) interphosphate distance, *d*_*b*_, (orange) and average unzipping force, *f*_m_ (black). Linear fits to data (black, blue, and orange lines, respectively) are also shown. (**C**) Example of application of the Clausius-Clapeyron relation to the experimental FDC (see text). (**D**) *T* -dependence of the ten NNBP entropies (red) and enthalpies (blue). Results are reported in Extended Data Tables 2 and 4, respectively. The entropy of motifs GC/CG and TA/AT were obtained by applying the circular symmetry relations. Fits to data (see text) are shown with a red (entropy) and blue (enthalpy) dashed line.

### T-dependence of the ssDNA elasticity

Deriving the full NNBP parameters in unzipping experiments requires measuring the *T* -dependent ssDNA elasticity. To do this, we extract the force versus the hairpin’s molecular extension (*x*) curve (hereafter referred to as FEC) with

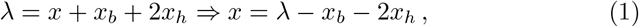

with *x* being shown as a green vertical line in Fig. 1A. Here, *x*_*b*_ and 2*x*_*h*_ are the bead displacement relative to the trap’s center and the handles extension, respectively (grey vertical lines in Fig. 1A). To determine the term *x*_*b*_ + 2*x*_*h*_ in Eq.(1), we have used the *effective stiffness* method ^48^ with *x*_*b*_ + 2*x*_*h*_ = *f/k*_eff_ and *x* = *λ − f/k*_eff_ where *k*_eff_ is obtained from a linear fit to the first slope in the FDC when the hairpin is fully folded (Sec. 1, Supp. Info). Notice that all extensions are *f* and *T* dependent.

From Fig. 1A, *x* = *x*_ss_ + *x*_*d*_, where *x*_ss_ is the ssDNA extension and *x*_*d*_ is the projection of the helix diameter (*d* = 2nm ^49^) on the pulling axis, which is described by Eq.(7) (Methods). To model the ssDNA elasticity, we use the intextensible WLC model ^50,51^ (Sec. 2, Methods). In this model, the extension *x*_ss_ at a given force is proportional to the number of released bases *n*, 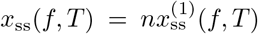 where *x*^(1)^ is the extension per base. For the fully folded hairpin, *n* = 0 and *x* = *x*_*d*_, whereas for the fully unzipped hairpin, 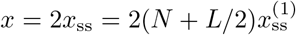 with *N* the number of base pairs in the stem and *L* the loop size. At a given *T*, we obtain 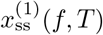 from the FEC measured after the last force rip (Extended Data Fig. 2). A fit of the WLC in Eq.(6) (Methods) to the rightmost part of the FEC gives the temperature-dependent persistence length (*l*_*p*_) and inter-phosphate distance (*d*_*b*_) of the ssDNA (Fig. 2A and Extended Data Table 1). As *T* increases, *l*_*p*_ (blue squares), varies from 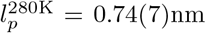 to 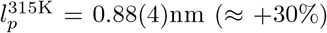. A linear fit to the data gives the slope 6(1) · 10^*−*3^nm/K (blue line). Moreover, the interphosphate distance, *d*_*b*_, (orange circles) shows a weak linear *T* -dependence (*≈* +5%) of slope 4(1) · 10^*−*4^nm/K (orange line). Similar behavior has been observed for shorter ssDNAs of 20 *−* 40 nucleotides ^52^ and polypeptide chains ^32^. The observed increase in *x*_ss_ with *T* is predicted in Debye-Huckel theory due to the entropy of the cloud of counterions. Upon increasing temperature, the screening of the phosphates repulsion is reduced, and *l*_*p*_ increases.

### Derivation of the NNBP entropies

To derive the entropies of the different NNBPs, we have decomposed the full unzipping curve into segments of variable length encompassing different regions along the FDC. Each segment is delimited by two peaks corresponding to force rips along the FEC. Figure 2C shows examples of segments starting and ending at a peak (colored circles). The entropy of hybridization, Δ*S*_0,*k*_(*T* ), of each segment *k* is given by (Sec. 3, Methods),

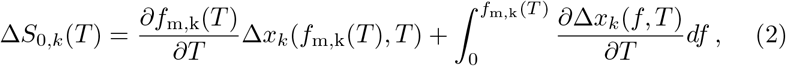

with Δ*x*_*k*_ the extension of segment *k*. Equation (2) is analogous to the Clausius-Clapeyron equation ^17,32,50^ in classical thermodynamics. Here *f* and *x* stand for the equivalent quantities of pressure and volume in hydrostatic systems. The r.h.s. of Eq.(2) depends on the average unzipping force of segment *k* measured at different temperatures, *f*_m,k_(*T* ), according to the equal area Maxwell construction for segment *k* (colored horizontal dashed lines in Fig. 2C). *f*_m,k_(*T* ) varies linearly with *T*, all segments showing the same slope *−*0.165(3)pN/K within statistical errors (Fig. 2B and Extended Data Table 1).

The integral in Eq.(2) accounts for the work needed to stretch the ssDNA and orient the molecule along the pulling axis between zero force and *f*_m,k_(*T* ) (Fig. 3A and Table 1, Extended Data). Equation (2) applied to the full FEC gives the total entropy of hybridization of the hairpin (Extended Data Fig. 3B).

**Table 1.**
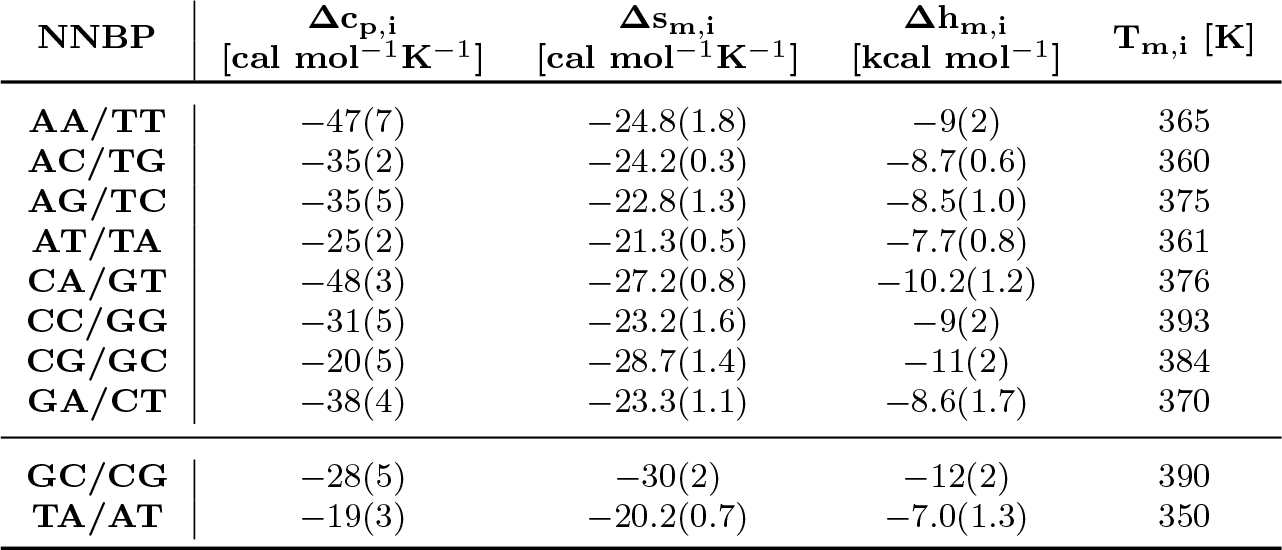
Measured heat capacity change (Δ*c*_*p,i*_), entropy (Δ*s*_*m,i*_), enthalpy (Δ*h*_*m,i*_), and melting temperature (*T*_*m,i*_) for the ten NNBP motifs. Errors are reported in brackets. Motifs GC/CG and TA/AT have been derived by applying the circular symmetry relations.

**Fig. 3.**
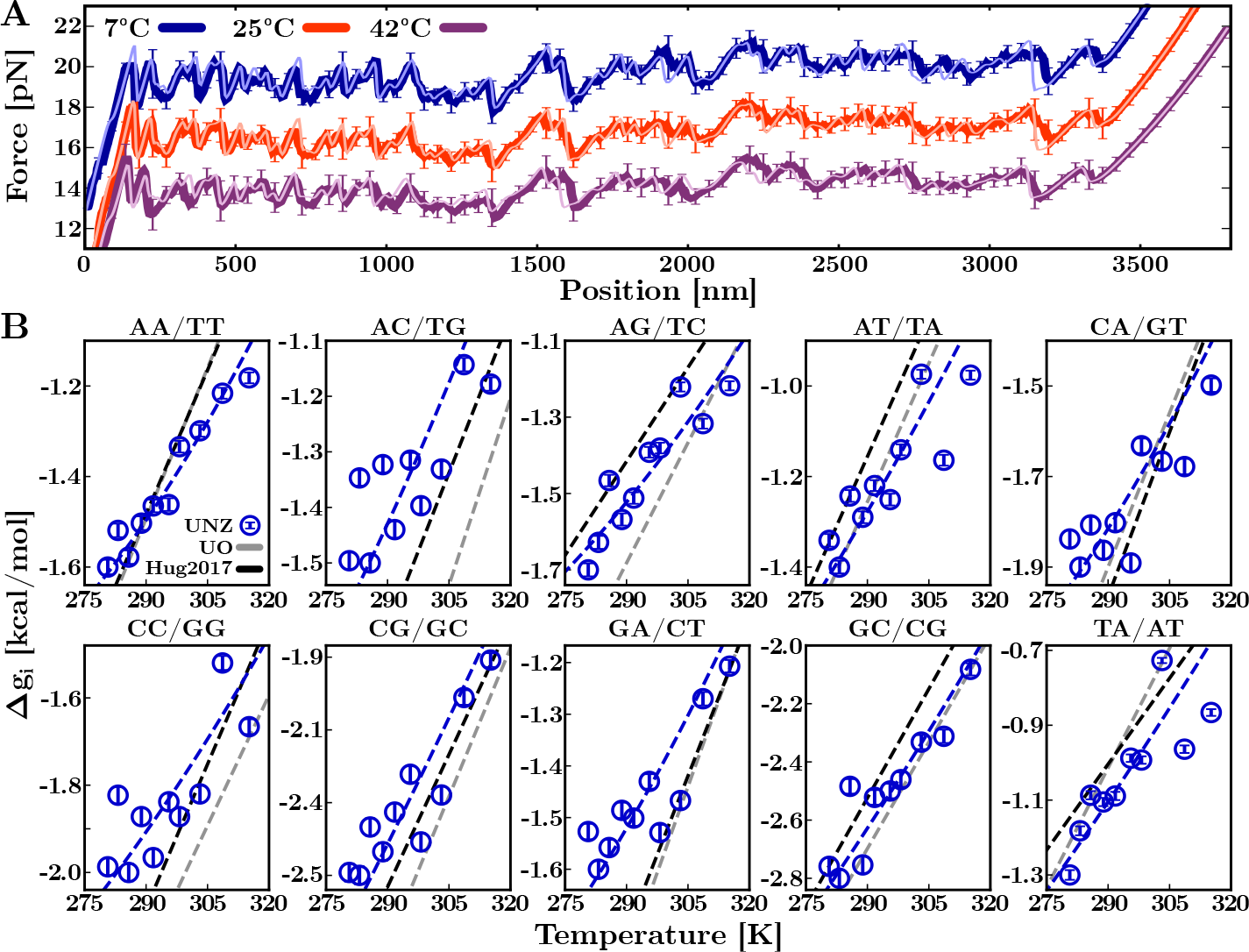
T-dependence of the NNBP DNA free energies. (**A**) Experimental FDCs (dark-colored lines) and theoretical predictions (light-colored lines) at 7^*°*^C, 25^*°*^C, and 42^*°*^C. Analogous results have been obtained at all temperatures (Extended Data Fig. 5). (**B**) Results for the ten NNBP DNA free energies (Extended Data Table 3). The free energy of motifs GC/CG and TA/AT has been computed with circular symmetry relations. A fit to data (blue line) has been added to compare with predictions by the UO (grey line) and Huguet *et al*. ^37^ (black line) sets.

To apply Eq.(2) for a given segment *k*, we must identify the DNA sequence limited by the initial and final peaks. A WLC curve passing through a peak at (*x, f* ) gives the number *n* of unzipped bases at that peak (dashed-grey lines in segment Δ*x*_*k*_). The initial and final values *n*_*A*_ and *n*_*B*_ (orange segment in Fig. 2C), identify the DNA sequence of that segment. Let *k* = 1, 2, …, *K* enumerate the different segments. The entropy of segment *k* at zero force and temperature *T* in the NN model is given by the sum of the individual entropies of all adjacent NNBPs within that segment,

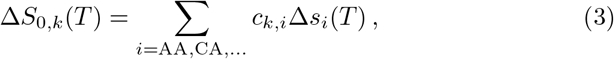

where the sum runs over the ten independent NNBP parameters labeled by the index *i*, and Δ*s*_*i*_ is the entropy of motif *i* with multiplicity *c*_*k,i*_, i.e., the number of times motif *i* appears in segment *k*. The entropy Δ*S*_0,*k*_(*T* ) in the l.h.s of Eq.(3) is measured using Eq.(2), and *c*_*k,i*_ for each motif is obtained from the segment sequence. A stochastic gradient descent algorithm has been designed to solve the system of *K* non-homogeneous linear equations (3) and derive the Δ*s*_*i*_ parameters at each *T* (Sec. 4, Methods). The results for the *T* -dependent DNA NNBP entropies Δ*s*_*i*_ are shown in Fig. 3D and reported in Extended Data Table 2. Typically, *K ∼* 400 *−* 600, making our single-molecule approach equivalent to melting experiments on different oligo sequences.

### NNBP free energies and enthalpies

From the previously derived NNBP entropies Δ*s*_*i*_(*T* ), we can also derive the NNBP enthalpies, Δ*h*_*i*_(*T* ), from the relation,

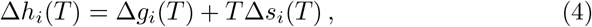

with Δ*g*_*i*_(*T* ) the free energies of the different motifs. To measure the Δ*g*_*i*_(*T* ), we have fitted the FDCs of Fig. 1B to the unzipping curves predicted by the NN model ^25,26,37^. The fitting procedure is based on a Monte-Carlo method that optimizes the eight independent energy parameters, Δ*g*_*i*_(*T* ), at each *T* (Sec. 5, Methods). The other two energy parameters (GC/CG and TA/AT) are obtained from the circular symmetry relations ^35–37^. Fits are shown in Fig. 3A for three selected temperatures, and the Δ*g*_*i*_(*T* ) are shown in Fig. 3B (see also Extended Data Table 3). Results (blue circles) agree with the unified oligonucleotide (UO) dataset (black line) and the energy parameters obtained by Huguet *et al*. in Ref. ^37^ (grey line). In this reference, unzipping experiments at room temperature (298K) were combined with melting temperature data of oligo hybridization over the vastly available literature. Overall agreement is observed, except for some motifs such as AC/TG and GA/CT where the UO energies are lower. The ten NNBP enthalpies were obtained from Eq.(4) at each *T* and are shown in Fig. 2D (see also Extended Data Table 4). The agreement between the new Δ*g*_*i*_(*T* ) values in Fig. 3B with previous measurements under the assumption that 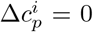 for all motifs ^37,43^, underlines the strong compensation between the temperature-dependent enthalpies and entropies shown in Fig. 2D that mask the finite 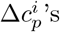.

### NNBPs heat capacity changes

The temperature-dependent NNBP entropies and enthalpies permit us to derive the heat capacity changes Δ*c*_*p,i*_ for all motifs by using the relations,

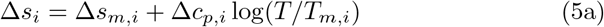

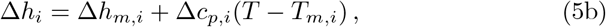

where *T*_*m,i*_ is the melting temperature of motif *i* where Δ*g*_*i*_(*T*_*m,i*_) = 0, and Δ*s*_*m,i*_ and Δ*h*_*m,i*_ are the entropy and enthalpy at *T*_*m,i*_, fulfilling Δ*h*_*m,i*_ = *T*_*m,i*_Δ*s*_*m,i*_. To derive the ten Δ*c*_*p,i*_, we fit the NNBP entropies to the equation *A*_*i*_ + Δ*c*_*p,i*_ log(*T* ), being *A*_*i*_ = Δ*s*_*m,i*_ *−* Δ*c*_*p,i*_ log(*T*_*m,i*_). The results are shown in Fig. 4A and Table 1 (column 1). From the Δ*c*_*p,i*_, we combined Eqs.(5) with Eq.(4) to fit the experimental values of Δ*g*_*i*_(*T* ) (blue dashed lines in Fig. 3B) to obtain *T*_*m,i*_. From the *T*_*m,i*_, we retrieve Δ*s*_*m,i*_ and Δ*h*_*m,i*_ from Eqs.(5a), (5b) (red and blue dashed lines in Fig. 2D). The fitting procedure is described in Sec. 4, Supp. Info. Results for *T*_*m,i*_, Δ*s*_*m,i*_ and Δ*h*_*m,i*_ are shown in Fig. 4B and Table 1. Notice the high *T*_*m*_ values of the individual motifs, a consequence of the high enthalpies of the NN motifs.

**Fig. 4.**
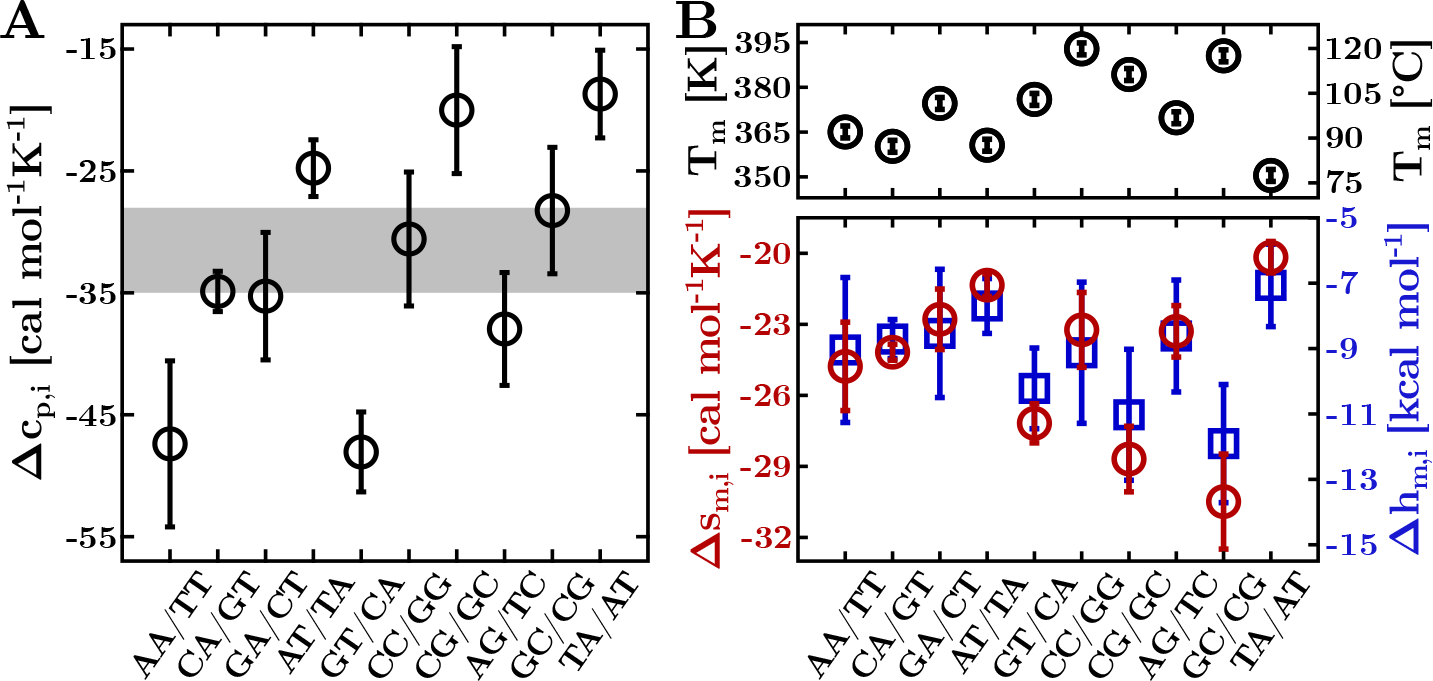
The DNA NNPB thermodynamics. (**A**) Measured heat capacity change per motif. The grey band shows the range of Δ*c*_*p*_ values per motif reported in Ref. ^12^. (**B**) Melting temperatures (top), entropies, and enthalpies at *T*_*m*_ (bottom) for each of the ten NNBP parameters. Results for motifs GC/CG and TA/AT have been derived by applying circular symmetry relations.

## Discussion

We measured the free energies, entropies, and enthalpies in the temperature range 7 *−* 42^*°*^C at the level of single nearest-neighbor base pairs (NNBP). We have mechanically unzipped a 3.6kbp DNA hairpin using a temperature-jump optical trap. The DNA sequence is long enough to permit us to accurately derive the ten NNBP free-energy parameters, Δ*g*_*i*_, by statistical modeling of the force-distance curve (FDC) ^26,37^. At first sight, the Δ*g*_*i*_ values (Fig. 3B) vary linearly with temperature due to the compensation of enthalpy and entropy in Eq.(4). This compensation masks the temperature dependence of enthalpies, Δ*h*_*i*_, and entropies, Δ*s*_*i*_, (cf. Eqs.(5)) arising from a finite Δ*C*_*p*_, rendering Δ*g*_*i*_ = Δ*h*_*i*_ *− T* Δ*s*_*i*_ linear in *T* . We have introduced an approach to derive the *T* -dependent entropies by combining the Clausius-Clapeyron relation in force (Eq.(2)) and the nearest-neighbor (NN) model for duplex formation. We implemented a tailored stochastic gradient descent algorithm to extract the ten *T* -dependent NNBP entropy parameters, Δ*s*_*i*_. Together with the Δ*g*_*i*_ values, the ten enthalpy parameters, Δ*h*_*i*_, readily follow. Fitting the results to Eqs.(5), we have obtained the Δ*c*_*p,i*_ and *T*_*m,i*_ values for the ten motifs (Fig. 4 and Table 1). Upon averaging over all motifs we get 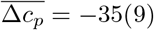 cal mol^*−*1^K^*−*1^bp^*−*1^. This must be compared to the highly dispersed results from bulk experiments ranging between -20 and -160 cal mol^*−*1^K^*−*1^bp^*−*1^, depending on the experimental technique, setup, and DNA sequence ^9,12^. In contrast, recent molecular dynamic simulations estimated an average Δ*c*_*p*_ *∼ −*30 cal mol^*−*1^K^*−*1^bp^*−*1 53,54^, in agreement with our results.

Force spectroscopy emerges as a reliable approach to accurately derive the energy parameters in NAs. Unzipping experiments control the unfolding reaction by moving the force-sensing device (e.g., optical trap in optical tweezers and cantilever in AFM). In DNA hairpins of a few kb, the sequence contains all ten NN motifs repeated several times, ensuring their reliable statistical sampling in single-DNA unzipping experiments. The high-temporal resolution combined with the sub-kcal/mol accuracy of work measurements permits us to derive the ten NN energy parameters at different temperatures. The main requirement of unzipping experiments is an accurate model of the elastic response of the single-stranded DNA (ssDNA). We have fitted the last part of the unzipping FDCs to the worm-like chain model, known to fit data well at high-salt conditions (1M NaCl) where ssDNA excluded-volume effects are negligible ^51^. Salt concentration might also affect the Δ*c*_*p,i*_ values. As these are related to the change in configurational entropy upon duplex formation, salt-dependent Δ*c*_*p*_’s might indicate a change in conformational heterogeneity in either the dissociated or hybridized strands upon varying salt.

How do our results compare to those derived from calorimetric melting experiments? The structure of the unfolded state differs in unzipping and thermal denaturation experiments, a fully stretched ssDNA at a given force, and a random coil at zero force, respectively. Their free energy difference equals the work to stretch the ssDNA from the random coil to the stretched conformation. Moreover, hybridization and unzipping differ in the order of the unfolding reactions: while hybridization of two complementary oligos, *A* and *Ā*, is a bimolecular reaction *A* + *Ā* ⇋ *AĀ*, hairpin unzipping is a unimolecular reaction *A* ⇋ *B* between the folded and unfolded conformations. Such difference is apparent in the dependence of *T*_*m*_ on the enthalpy 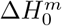 and entropy 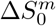 of folding. For a bimolecular reaction, the value of *T*_*m*_ explicitly depends on the total oligo concentration *c*, with 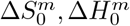 taken at the reference 1M salt condition (Eq.(10), Methods). Instead, for the unimolecular unzipping reaction, 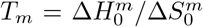 does not include the entropy of mixing the dissociated strands. We expect that temperature-dependent enthalpies Δ*h*_*i*_ are equal for hybridization and unzipping, whereas entropies Δ*s*_*i*_ should differ due to the entropy of mixing. We have determined the homogeneous entropy correction *δ*Δ*s* to the total entropy 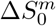 between hybridization (bimolecular) and unzipping (unimolecular) reactions (Sec. 6, Methods). The correction is an intensive quantity that is independent of oligo sequence and length, *δ*Δ*s* = 6(1) cal mol^*−*1^K^*−*1^ *∼* 4*R* log 2, where *R* = 1.987 cal mol^*−*1^K^*−*1^ is the ideal gas constant (Eq.(11), Methods). This value has been obtained by comparing the *T*_*m*_ values predicted by our energy parameters using Eq.(10) with the experimental values obtained for DNA duplexes in Ref. ^55^ at 1020mM NaCl and *c* = 2*µ*M. The results of such a comparison are shown in Extended Data Fig. 6. Practically, the effect of the entropic correction *δ*Δ*s* on *T*_*m*_ is small as it is *∼* 3 *−* 4 times lower than the average NNBP’s entropy 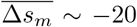 cal mol^*−*1^K^*−*1^ (cf. Table 1). Notice that 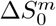 is extensive, growing linearly with the oligo length, whereas *δ*Δ*s* is intensive. Therefore, the correction *δ*Δ*s ∼* 5 cal mol^*−*1^K^*−*1^ is negligible for sufficiently large oligos, being already 5% for oligos of just ten nucleotides and further decreasing for longer DNAs (see Extended Data Table 5).

## Conclusions

The remarkable accuracy of the nearest-neighbor model for reproducing the experimentally measured force-distance curves permitted us to derive the temperature-dependent DNA energy parameters. One might ask whether there are deviations from the NN model, e.g., in the form of next-to-nearest neighbor (NNN) effects predicted to be important for some tetranucleotide motifs ^56^. However, NNN effects might be difficult to observe in unzipping experiments of long DNA hairpins. Our method averages local effects over the whole sequence hiding potential deviations from the NN model at some locations. Unzipping studies on suitably designed short DNA hairpins containing specific NNN motifs would be more appropriate to address this problem. In this case, determining the elastic response of the specific ssDNA sequence would also be necessary ^57^. The unzipping method might also be applicable to derive the temperature-dependent energy parameters of RNA, where finite Δ*C*_*p*_ effects are particularly relevant to the RNA folding problem. Previous studies at room temperature show that RNA unzipping is an irreversible process driven by stem-loops forming along the unpaired strands that compete with the hybridization of the native stem ^44^. Such irreversible and kinetic effects suggest higher Δ*c*_*p,i*_ values in RNA compared to DNA. DNA thermodynamics down to 0^*°*^C might find applications for predicting DNA thermodynamics at low temperatures and cold denaturation effects. Estimates based on our Δ*c*_*p,i*_ values show that the most stable DNA hairpins ending a tetraloop predict cold denaturation temperatures lower than *≈ −*90^*°*^C (Extended Data Fig. 7), raising questions about the importance of cold denaturation effects for cryophile organisms surviving in extremely cold environments ^58^. Finally, our results have implications for determining the DNA force fields used in molecular dynamics simulations of DNA conformational kinetics ^59^, essential for computational studies of NAs in general.

## Methods

### 1 Molecular Construct and Experimental Setup

We used a temperature-jump optical trap ^17^ (OT) to unzip a 3593bp DNA hairpin flanked by short (29bp) DNA handles and ending with a GAAA tetraloop. Experiments have been carried out in the temperature range [7,42]°C in a buffer of 1M NaCl, 10mM Tris-HCl (pH 7.5), and 1mM EDTA. Experiments have been performed at a constant pulling speed, *v* = 100nm/s. We sampled 5-6 different molecules at each temperature, collecting a minimum of *∼* 50 unfolding-folding trajectories per molecule. To change the temperature inside the microfluidics chamber, the MiniTweezers setup implements a heating laser of wavelength *λ* = 1435nm. This device allows for increasing the temperature by discrete amounts of Δ*T ∼* +2.5^*°*^C up to a maximum of *∼* +30^*°*^C with respect to the environment temperature, *T*_0_. By placing the OT in an icebox cooled down at a constant *T*_0_ *∼* 5^*°*^C, it has been possible to carry out experiments at a minimum temperature of 7^*°*^C. The design of the microfluidics chamber has been chosen to damp convection effects caused by the laser non-uniform temperature, which may produce a hydrodynamics flow between medium regions (water) at different *T* .

In a typical OT unzipping experiment, the molecule is tethered between two polystyrene beads through specific interactions with the molecular ends. One end is labeled with a digoxigenin (DIG) tail and binds with an anti-DIG coated polystyrene bead (AD) of diameter 3*µ*m. The other molecular end is labeled with biotin (BIO) and binds with a streptavidin-coated bead (SA) of diameter 2*µ*m. The SA bead is immobilized by air suction at the tip of a glass micropipette, while the AD bead is optically trapped. A pulling cycle consists of moving the optical trap between two fixed positions: the molecule starts in the folded state, and the trap-pipette distance (*λ*) is increased, resulting in an external force to be applied to the molecular ends. This causes the hairpin to progressively unzip in a stick-slip process characterized by a sequence of slopes and force rips (FDC sawtooth pattern). When the molecule is completely unzipped, the rezipping protocol starts; *λ* is decreased, and the molecule folds back reversibly to the hairpin (native) state.

### 2 ssDNA elastic model

The total hairpin extension with *n* unzipped bases at force *f* and temperature *T, x*(*n, f, T* ), is given by the sum of the ssDNA extension (*x*_ss_(*f, n, T* )) plus the contribution of the double helix diameter (*x*_*d*_(*f, T* )). The ssDNA elastic response has been modeled according to the Worm-Like chain (WLC), which reads

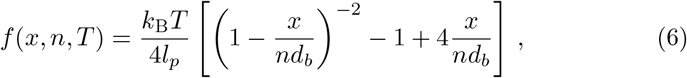

where *k*_B_ is the Boltzmann constant, and *T* is the temperature, *l*_*p*_ is the persistence length, *d*_*b*_ is the interphosphate distance and *n* is the number of ssDNA bases. Notice that the computation of the ssDNA extension requires inverting Eq.(6) ^48^.

The observed increase in the ssDNA extension with *T* at a given force (Fig. 1B) demonstrates that *l*_*p*_ and *d*_*b*_ are *T* -dependent. This contrasts with the original assumption in the WLC model that *l*_*p*_ and *d*_*b*_ are temperature independent and *x*_ss_ decreasing with *T* at a given force. Remarkably, Eq.(6) accurately describes our data as stacking effects in mixed purine-pyrimidine sequences are negligible in our experimental conditions ^51^.

The contribution of the hairpin diameter to *x* is given by the projection of the helix diameter in the direction of propagation of the force. It is described as a free dipole in an external force field and is modeled by the FJC model, which reads

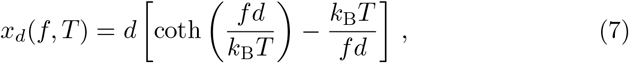

where *d* = 2nm is the hairpin diameter ^49^.

### 3 The Clausius-Clapeyron Equation

In the unzipping experiment, the total trap-pipette distance is given by *λ* = *x*+*x*_*b*_+2*x*_*h*_, where *x* is the extension of the ssDNA plus the molecular diameter, while *x*_*b*_ and 2*x*_*h*_ are the bead displacement relative to the trap’s center and the handles extension, respectively. Starting with the hairpin totally folded, unzipping consists in converting the double-stranded DNA into ssDNA until it is completely unfolded. The free energy needed to fold back the ssDNA into the hairpin and decrease the applied force to zero is given by

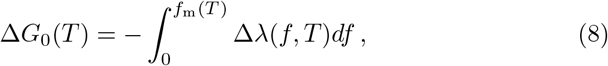

where Δ*λ* is the total extension change of the hairpin between the initial and final states of the unzipping integrated between zero and the mean unzipping force, *f*_*m*_. Notice that the contributions from *x*_*b*_ and 2*x*_*h*_ to *λ* remain constant during the entire unzipping process so that Δ*λ* = Δ*x*, i.e., equals the extension change due to the ssDNA and the molecular diameter only.

The folding entropy can be directly derived from the thermodynamic relation Δ*S*_0_ = *−∂*Δ*G*_0_*/∂T*, which gives

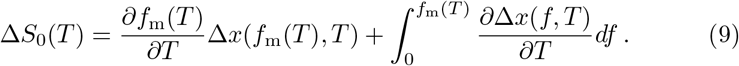

The first term of Eq.(9) is analogous to the Clausius-Clapeyron equation for first-order phase transitions, where *f* and *x* are equivalent to pressure and volume. The integral term in Eq.(9) accounts for the (positive) entropic contribution to stretch the ssDNA and orient the molecular diameter along the pulling axis from zero to force *f*_*m*_.

### 4 Stochastic Gradient Descent Method

The derivation of the DNA NNBP entropies corresponds to solving the non-homogeneous linear system of *K* equations and *I* = 8 parameters given by Eq.(3). To do this, we used a custom-designed stochastic gradient descent (SGD) algorithm. The model is described in Sec. 2, Supp. Info. The application of an optimization algorithm to this problem is made possible by the large data set built accounting for all possible peaks combination, which gives *K ∼* 400 *−* 600 *≫ I* for each experimental *T* .

### 5 Derivation of the NNBP Free-Energies

Starting with an initial guess of the ten independent Δ*g*_*i*_, a random increment of the energies is proposed at each optimization step, and a prediction of the FDC is generated. The latter is given by the competition of two energy contributions at each position of the optical trap (*x*_tot_): the energy of the stretched molecular construct acting to unfold the molecule (Δ*G*_el_(*x*_tot_)), and the energy of the hybridized bps keeping the hairpin folded (Δ*G*_0_(*x*_tot_, *n*)). At a given *x*_tot_ and *n*_1_ hybridized bp (Fig. 2A), the hairpin unfolds when Δ*G*_el_(*x*_tot_) *>* Δ*G*_0_(*x*_tot_, *n*_1_) (force rip) and Δ*n* = *n*_2_ *− n*_1_ bp are released lowering the stretching contribution so that Δ*G*_el_(*x*_tot_) *<* Δ*G*_0_(*x*_tot_, *n*_2_) (see Sec. 3, Supp. Info). The error in approximating the experimental FDC with the theoretical one, *E* = (*f*_exp_ *− f*_theo_)^2^, drives a Metropolis algorithm: a change of the energy parameters is accepted if the error difference to the previous step is negative (Δ*E <* 0). Otherwise (Δ*E >* 0), the proposal is accepted if exp(*−*Δ*E/T* ) *< r* with *r* a random number uniformly distributed *r ∈ U* (0, 1). The algorithm continues until convergence (until Δ*E* is smaller than a given threshold) or until the maximum number of iterations is reached. The parameters corresponding to the smallest value of *E* are the optimal NNBP free energies.

### 6 Prediction of the Oligos Melting Temperature

The melting temperature for non-self-complementary sequences in bimolecular reactions (hybridization) is given by

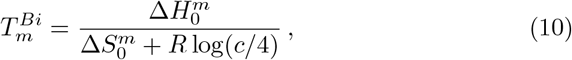

where 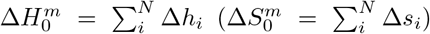 is the total duplex enthalpy (entropy) at *T* = *T*_*m*_, *R* = 0.001987 kcal mol^*−*1^K^*−*1^ is the ideal gas constant, *c* is the experimental oligo concentration. In contrast, for unimolecular reactions (folding), we found

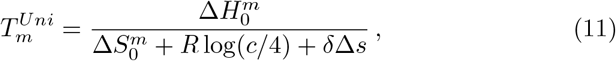

where *δ*Δ*s* = 4*R* log 2 is a correction to the total entropy 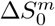. By subtracting the inverse of Eq.(10) and Eq.(11), one gets

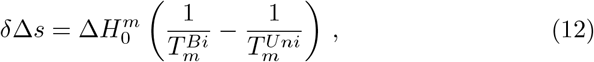

where the term in parenthesis is computed by subtracting the *T*_*m*_ values predicted by our energy parameters using Eq.(10) to the experimental dataset measured by Ocwzarzy *et al*. at 1020mM NaCl and *c* = 2*µ*M ^55^ (Extended Data Fig. 6 and Table 5). Finally, 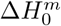 is determined by using our NNBP enthalpy parameters.

## Supplementary information

This article has accompanying Supplementary Information and Extended Data.

## Acknowledgments

P.R. was supported by the Angelo Della Riccia Foundation. M.R.-P. was supported by the Spanish Research Council JDC2022-049996-I. F.R. was supported by Spanish Research Council Grant PID2019-111148GB-I00 and the Institució Catalana de Recerca i Estudis Avançats (F. R., Academia Prize 2018).

## Authors’ contributions

P.R. and M.R.-P. carried out the experiments. S.B.S. designed and built the instrument. P.R. and M.R.-P. analyzed the data. P.R., M.R.-P., and F.R. designed the research. P.R. and F.R. wrote the paper.

