## Supplementary methods for "DNA Calorimetric Force Spectroscopy at Single Base Pair Resolution"

### 1 Temperature Dependence of the DNA FDCs

The elastic properties of ssDNA are strongly temperature dependent (see Fig. 1B, main text). Accurately measuring these properties requires modeling all contributions to the trap-pipette distance, $\lambda$, which includes the optical trap displacement ($x_{\rm b}$), the dsDNA handles ($x_{\rm h}$), the ssDNA ($x_{\rm ss}$), and the molecular diameter ($d_0$). The setup contributions ($x_{\rm b}$ and $x_{\rm h}$) can be evaluated by using the *effective stiffness* method[48]. According to it, these terms are approximated by an effective stiffness, $k_{\rm eff}^{-1} \approx k_{\rm h}^{-1} + k_{\rm b}^{-1}$. The use of short handles (29bp) makes the evaluation of the stretching terms easier as their stiffness is much larger as compared to the trap stiffness ($k_{\rm h} \gg k_{\rm b}$), implying that $k_{\rm eff} \approx k_{\rm b}$. Moreover, if the force varies in a relatively narrow range ($f_{\rm max} - f_{\rm min} \lesssim$ 10pN), the trap stiffness can be considered nearly force-independent so $k_{\rm eff}$ is constant along the folded branch of the FDC. Therefore, we can estimate $k_{\rm eff}$ by fitting the slope preceding the first force rip in the FDC to the linear equation $f = k_{\rm eff} x$ (orange dashed-line in Extended Data Fig. 1). This allows us to compute the (effective) contribution of the handles and optical trap, $x_{\rm eff}$, to the total trap-pipette distance, $\lambda$.

### 2 Stochastic Gradient Descent in a Nutshell

The basic principle behind stochastic approximation can be backtracked to the Robbins–Monro algorithm of the 1950$s$[60]. Since then, stochastic gradient descent (SGD) methods have become one of the most widely used optimization methods[61–65]. SGD is an iterative method for optimizing an objective function, $J(w)$, with suitable smoothness properties (e.g., differentiable or subdifferentiable). The set of parameters, $w^*$, minimizing $J(w)$, is iteratively approximated according to an update algorithm proportional to the antigradient of the objective function, $-\nabla_w J(w)$. Starting from an initial guess of $w$, at each step of the algorithm, the parameters are updated according to

$$\begin{cases} w_{t+1} & = w_t + v_{t+1} \\ v_{t+1} & = \beta v_t - \eta \nabla_{w_t} J(w)\,, \end{cases} \tag{1}$$

where $v_t$ is the *velocity* of the optimization and $\eta \geq 0$ is the step size (called *learing rate*). The parameter $\beta$ (the so-called *momentum coefficient*) accounts for a fraction of the previous step in the current update. The critical difference between SGD and standard gradient descent algorithms is that information (total entropy and coefficients) from only one FEC segment ($\Delta x_k$) at a time is used to calculate the step, and the segment is picked randomly at each step.

The SGD convergence rate can be improved by considering Nesterov's Accelerated Gradient (NAG), introduced in 1983[66,67]. According to NAG, the update equations are

$$\begin{cases} w_{t+1} & = w_t + v_{t+1} \\ v_{t+1} & = \beta v_t - \eta \nabla_{w_t+\beta v_t} J(w)\,. \end{cases} \tag{2}$$

While the classic momentum (CM) algorithm updates the velocity vector by computing the gradient at $w_t$, the NAG algorithm computes the gradient at $w_t + \beta v_t$. To make an analogy, while CM faithfully trusts the gradient at the current iteration, NAG puts less faith in it and looks ahead in the direction suggested by the velocity vector; it then moves in the direction of the gradient at the look-ahead point. If $\nabla_{w_t+\beta v_t} J(w) \approx \nabla_{w_t} J(w)$, then the two updates are similar. The advantage of using NAG is that it converges at a rate of $\mathcal{O}(1/t^2)$, while CM converges at a rate of $\mathcal{O}(1/t)$.

To derive the DNA NNBP entropies from unzipping experiments, we used an SGD minimization implementing NAG update equations. Let us rewrite Eq.(2) (main text) as $\mathbf{\Delta S_0} = C\mathbf{\Delta s}$, where $\mathbf{\Delta S_0}$ is the vector of entropies measured with the Clausius-Clapeyron equation for each of the $K$ FEC segments, $\mathbf{\Delta s}$ is the vector of the $I = 8$ NNBP entropy parameters, and $C$ is the $K \times I$ matrix of the coefficients, $c_{k,i}$.

Thus, for a given loss function (ex., least squares), the algorithm has to minimize

$$J(w) = \sum_{k=1}^{K} (\hat{w}_k - w_k)^2 = \sum_{k=1}^{K} (\Delta S_{0,k} - C_k \mathbf{\Delta s})^2\,. \tag{3}$$

By using this method, we measured the DNA entropies at the single base-pair level for each experimental temperature in the range $[280, 315]$ K (see results in Fig. 3C, main text and Extended Data Table 3).

### 3 Prediction of the DNA Unzipping Curve

In unzipping experiments, the total trap–pipette distance, $\lambda$, can be written as

$$\lambda(f) = x_{\mathrm{b}}(f) + x_{\mathrm{h}}(f) + x_{\mathrm{ss}}(f, n) + x_{\mathrm{d}}(f)\,, \tag{4}$$

where $x_{\mathrm{b}}(f)$ is the displacement of the bead from the center of the optical trap, $x_{\mathrm{h}}(f)$ and $x_{\mathrm{ss}}(f, n)$ account for the extension of the two double-stranded handles and the ssDNA extension, respectively (described with the WLC model, Eq.(5), Methods), and $x_{\mathrm{d}}(f)$ is the projection of the folded hairpin of diameter $d$ (typically $d = 2$nm for DNA and RNA hairpins [49]) along the pulling axis [68]. It is modeled with the freely-jointed chain in Eq.(6), Methods. For a given $\lambda$, the total system free energy is given by

$$\begin{aligned} \Delta G_{\mathrm{tot}}(\lambda, n) =& \Delta G_0(n) + \Delta G_{\mathrm{b}}(x_{\mathrm{b}}) + \Delta G_{\mathrm{h}}(x_{\mathrm{h}}) + \\ &+ \Delta G_{\mathrm{ss}}(x_{\mathrm{ss}}, n) + \Delta G_{\mathrm{d}}(x_{\mathrm{d}})\,, \end{aligned} \tag{5}$$

where $\Delta G_0(n) = \sum_i^n \Delta g_{0,i}$, is the hairpin free-energy of hybridization according to the NN model and the other terms are the energy contributions of the corresponding elastic terms in Eq.(4)

### 3.1 Computation of the Equilibrium FDC

Let us consider the case where thermal fluctuations are neglected in the FDC computation. Thus, at a given value of $\lambda$, the system is always in the state of minimum energy, $\Delta G_{\mathrm{eq}}(\lambda) = \Delta G_{\mathrm{tot}}(\lambda, n^*)$. To compute the equilibrium free energy of the system, let us first introduce the system partition function, $Z$. At each $\lambda$, this is defined as the sum over all the possible states, i.e., all the possible sequences of $n$ open base pairs, which is

$$Z(\lambda) = \sum_{n=0}^{N} \exp\left(-\frac{\Delta G_{\mathrm{tot}}(\lambda, n)}{k_{\mathrm{B}}T}\right), \tag{6}$$

where $N$ is the total number of base pairs of the sequence. Finally, by recalling that $\Delta G = -k_{\mathrm{B}}T \ln Z$, the equilibrium force is given by:

$$f_{\mathrm{eq}}(\lambda) \equiv \frac{\partial \Delta G(x_{\mathrm{eq}})}{\partial \lambda} = -k_{\mathrm{B}}T \frac{\partial \ln Z(\lambda)}{\partial \lambda}. \tag{7}$$

Computing Eq.(6) requires solving the transcendental Eq.(4) with respect to $f$ (that can be performed numerically) and then computing Eq.(5) for all $n \in [0, N-1]$. For each $\lambda$, the value $n^*$ minimizing the equilibrium free-energy $\Delta G_{\mathrm{eq}} = \Delta G_{\mathrm{tot}}(\lambda, n^*(\lambda))$ gives the most probable number of open base-pairs. Eventually, the computation of the equilibrium force in Eq.(7) gives a theoretical prediction for the unzipping curve of a given sequence (Extended Data Fig. 5).

### 3.2 Equilibrium Free Energy

The total free energy in Eq.(5) is the sum of two main contributions: the hybridization energy $\Delta G_0(n)$, which linearly depends on the number of hybridized NNBPs $n$, and the stretching energy $\Delta G_{\mathrm{el}}(\lambda, n) = \Delta G_{\mathrm{b}}(x_{\mathrm{b}}) + \Delta G_{\mathrm{h}}(x_{\mathrm{h}}) + \Delta G_{\mathrm{ss}}(x_{\mathrm{ss}}, n) + \Delta G_{\mathrm{d}}(x_{\mathrm{d}})$ depending on both $n$ and $\lambda$. For a given $\lambda$, the equilibrium configuration of the system is that with minimum $\Delta G_{\mathrm{el}}(\lambda, n^*)$ and maximum $\Delta G_0(n^*)$ among all possible values of $n$. Notice that for a hairpin of $N$ bp, $n$ ranges from 0 (native state) to $N-1$ NNBPs (totally unfolded), which gives $N-1$ possible system configurations for each value of $\lambda$.

Let us suppose that the system starts at equilibrium, with $n_1$ open bp. Upon increasing $\lambda$, the elastic term in Eq.(5) also increases. The number of open bp, $n_1$, remains constant until a value of $n = n_2 > n_1$ is found so that $\Delta G_{\mathrm{tot}}(\lambda, n_1) \equiv \Delta G_{\mathrm{tot}}(\lambda, n_2)$ (Extended Data Fig. 4A, top): even though the total energy of these two states is the same, the energetic internal balance is different (Extended Data Fig. 4A, bottom). The system minimizes the elastic free energy and switches to state $n_2$ by releasing $\Delta n = n_2 - n_1$ bp. Notice that, despite opening $\Delta n$ bp increases the system's energy, the released ssDNA causes the elastic contribution to decrease. In general, $\Delta G_{\rm el} \gg \Delta G_0$ so the global balance of the state $n_2$ is lower than the one of $n_1$. Therefore, the equilibrium free energy of hybridization, $\Delta G_0(n*)$, is a step function increasing with $\lambda$ (Extended Data Fig. 4B) with each discontinuity corresponding to a rip along the equilibrium FDC.

### 4 Fit of the NNBP parameters

The $T$-dependent NNBP entropies and enthalpies permit us to derive the heat capacity changes $\Delta c_{p,i}$ for each motif from the relations,

$$\Delta s_i = \Delta s_{m,i} + \Delta c_{p,i} \log(T/T_{m,i}) \tag{8a}$$

$$\Delta h_i = \Delta h_{m,i} + \Delta c_{p,i}(T - T_{m,i})\,, \tag{8b}$$

where $T_{m,i}$ is the melting temperature of motif $i$, and $\Delta s_{m,i}$ and $\Delta h_{m,i}$ are the entropy and enthalpy at $T = T_{m,i}$, respectively. The extraction of the NNBP thermodynamics parameters $(\Delta c_{p,i}, \Delta s_i, \Delta h_i, T_{m,i})$ has to be carried out carefully as the results are susceptible to experimental errors and parameters initialization. In particular, $\Delta s_{m,i}$, $\Delta h_{m,i}$, and $T_{r,i}$ strongly depend on their initialization values when directly fitted from Eqs.(8) as an error in $\Delta s_{m,i}$ $(\Delta h_{m,i})$ get compensated by $T_{m,i}$ and *vice versa*.

To derive the $\Delta c_{p,i}$, we fit the NNBP entropies to the equation $\Delta s_i(T) = A_i + \Delta c_{p,i} \log(T)$, being $A_i = \Delta s_{m,i} - \Delta c_{p,i} \log(T_{m,i})$. Notice that we derive $\Delta c_{p,i}$ from the NNBP entropies as they are obtained from the experimental data, in contrast to enthalpies that are computed from the free energies. Given $\Delta c_{p,i}$, we fit the NNBP free energies, $\Delta g_i(T)$, to the equation

$$\begin{aligned}\Delta g_i(T) &= \Delta h_i(T) - T\Delta s_i(T) = \\ &= \Delta h_{m,i} + \Delta c_{p,i}(T - T_{m,i}) - T\left(\Delta s_{m,i} + \Delta c_{p,i} \log\left(\frac{T}{T_{m,i}}\right)\right) \\ &= B_i + \Delta c_{p,i} T - T\left(A_i + \Delta c_{p,i} \log\left(T\right)\right)\,.\end{aligned} \tag{9}$$

obtained by combining Eqs.(8) (blue dashed lines in Fig. 3B, main text). Notice that $B_i = \Delta h_{m,i} - \Delta c_{p,i} T_{m,i}$. By definition, $T_{m,i}$ is the high temperature value where $\Delta g_i(T_{m,i}) = 0$. Finally, a new fit to Eqs.(8a) and (8b) by using the previously derived values of $\Delta c_{p,i}$ and $T_{m,i}$ (red and blue dashed lines in Fig. 2D, main text), gives $\Delta s_{m,i}$ and $\Delta h_{m,i}$. The results are shown in Fig. 4 and Table 1 of the main text.
