## Supplementary Figures and Tables for "DNA Calorimetric Force Spectroscopy at Single Base Pair Resolution"

#### Contents

**Extended Data: Figures** **2**

**Extended Data: Tables** **9**

### Extended Data: Figures

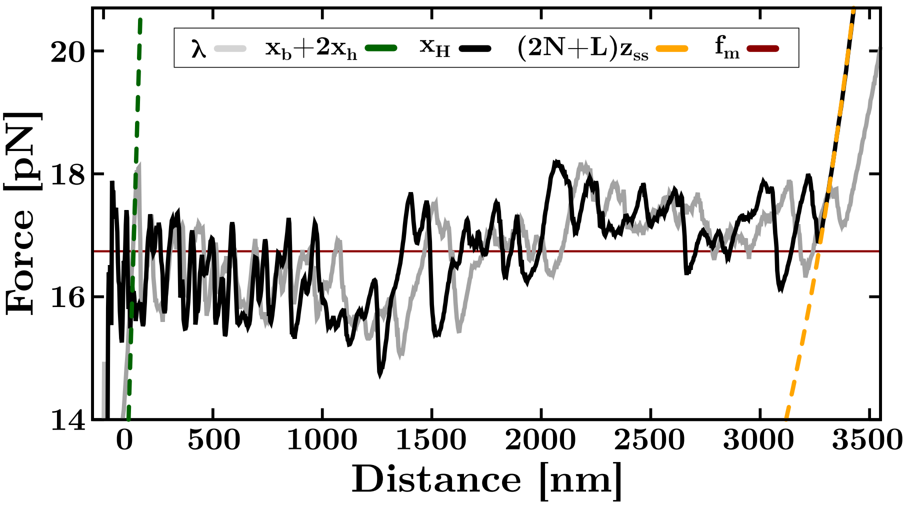


**Fig. 1**: **Computation of the FEC from the experimental FDC**. The force versus hairpin extension, $x_H$, (black line) is computed by subtracting to the trap position, $\lambda$, (grey line) the elastic contribution of the optically trapped bead, $x_b$, and DNA handles, $2x_h$, (green dashed line). To measure the $T$-dependent ssDNA elasticity, we fit the FEC after the last rip to the WLC model (orange dashed line). Notice that the average unzipping force, $f_m$, (red line) remains constant upon computing the FEC. Data are shown at $T = 25°$C.

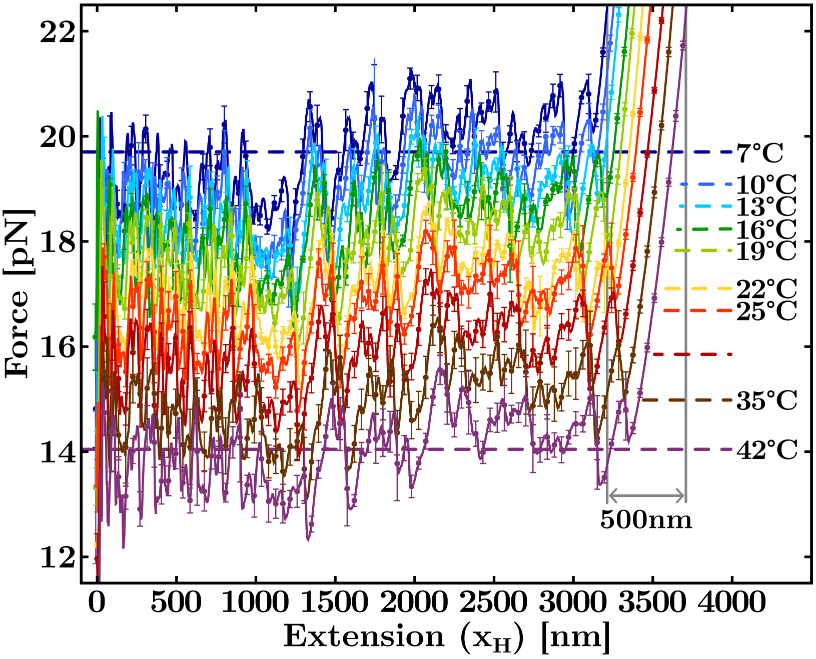


**Fig. 2**: **T-dependence of the measured force versus hairpin extension.** At each $T$, average unzipping forces are shown by dashed lines. The extension change over the studied temperature range is $\sim$ 500nm (grey vertical lines).

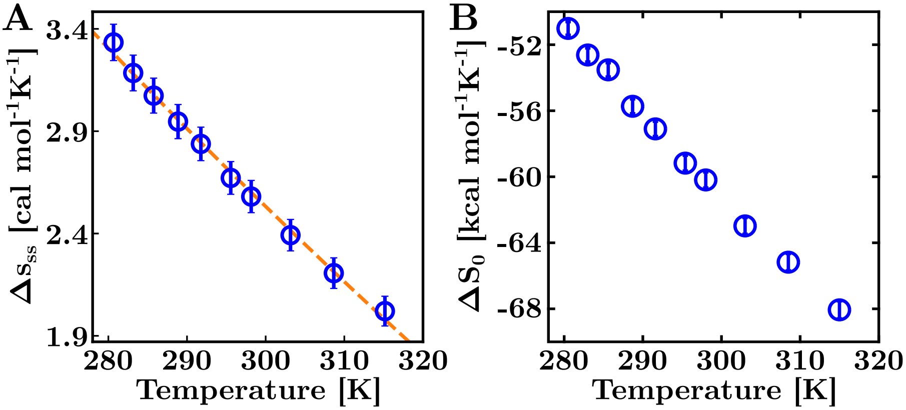


**Fig. 3**: **Clausius-Clapeyron equation applied over the full FDC.** (**A**) $T$-dependence of the entropy change per base, $\Delta s_{ss}$. It accounts for the work to stretch the ssDNA and orient the folded molecule along the direction of the external force, from $f = 0$pN to $f_{\mathrm{m}}(T)$ (integral term in Eq.(2), main text). The results are reported in Extended Data Table 1. A fit to data according to $\Delta s_{\mathrm{ss}}(T) = \Delta s_{\mathrm{ss},0} + \Delta c_p^{\mathrm{ss}} \log(T/T_m)$ (orange dashed line), gives the ssDNA heat capacity change per base at zero force, $\Delta c_p^{\mathrm{ss}} = -11.2 \pm 0.2\,\mathrm{cal\,mol^{-1}K^{-1}}$. (**B**) $T$-dependence of the total entropy change, $\Delta S_0(T)$, upon unzipping the 3.6kbp DNA hairpin measured using the Clausius-Clapeyron equation (see Eq.(2), main text).

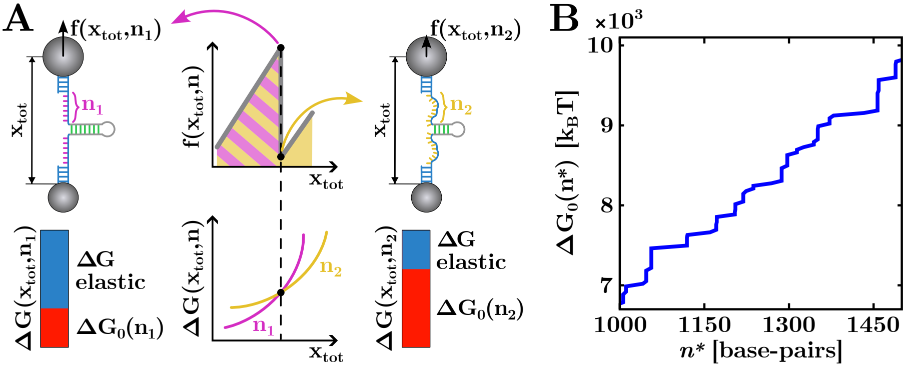


**Fig. 4**: **Derivation of the theoretical FDC.** (**A**) Schematics of the stretching and hybridization energy contributions. Upon unzipping, the molecule has $n_1$ open bp before the force rip (left) and $n_2 > n_1$ open bp after the rip (right). At the force rip (black dots), the total free energy of the system is the same in both states, and the system changes from the highest free energy branch ($n_1$, pink) to the lowest energy branch ($n_2$, pink). (**B**) The free energy of hybridization upon unzipping the hairpin is a monotonically increasing step function, with each discontinuity corresponding to a rip along the equilibrium FDC.

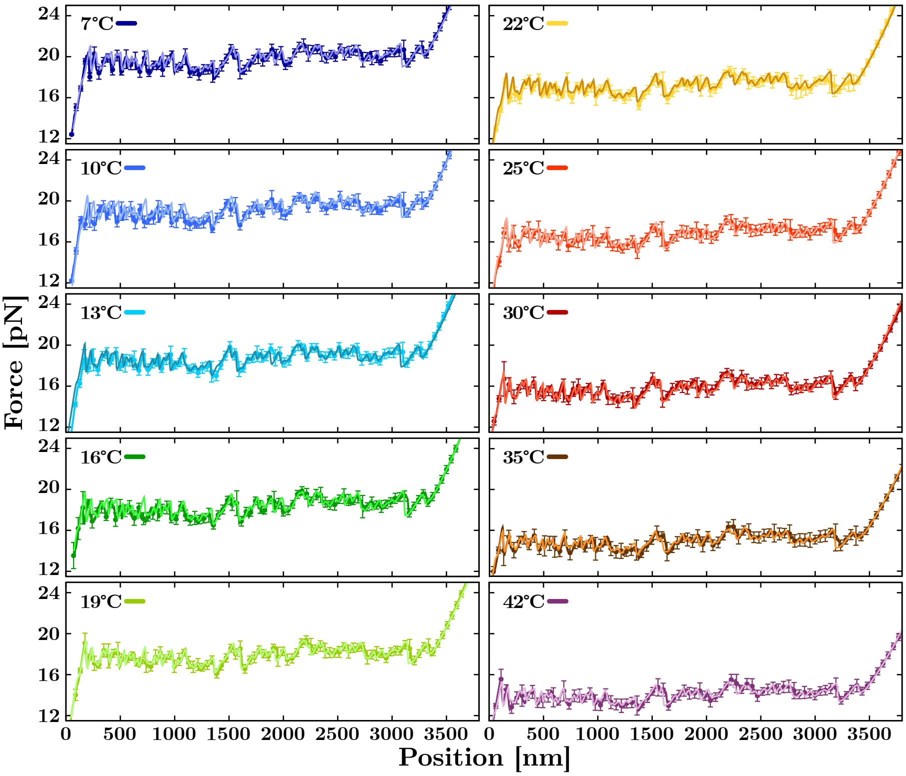


**Fig. 5**: **T-dependent theoretical FDC predictions.** Experimental FDCs (dark-colored solid lines and points) and theoretical predictions (light-colored lines) obtained with the free energy parameters derived at each $T$. Error bars indicate the variability of the experimental FDCs.

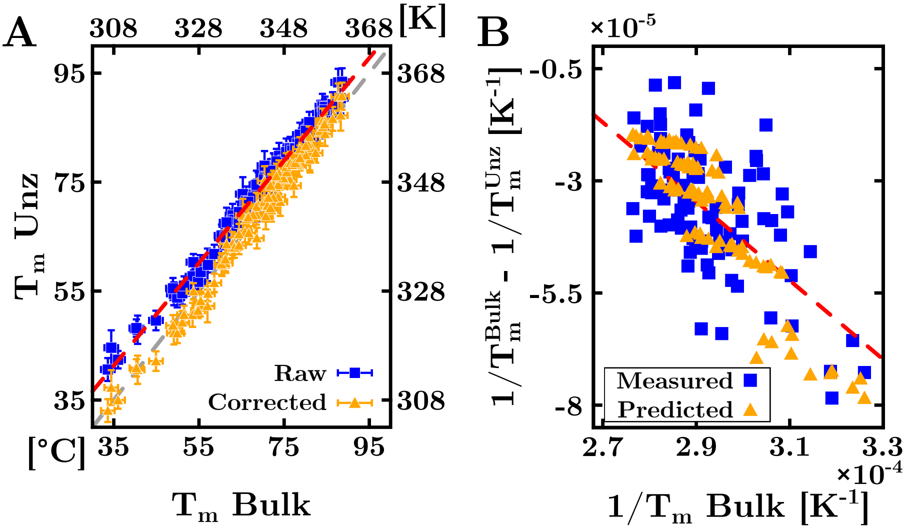


**Fig. 6**: **Prediction of the DNA duplexes melting temperatures.** (**A**) Comparison of the melting temperatures for the set of 92 DNA oligos studied by Owczarzy *et al.* in Ref.[55] (horizontal axis) and the values predicted with the unzipping energy parameters (vertical axis). Perfect agreement between the two data sets would imply all points falling on the dashed grey line $x = y$. Predictions obtained with Eq.(10) of Sec. 6, Methods (blue squares) show a systematic discrepancy with respect to the experimental values (dashed red line). By accounting for the entropic correction (Eq.(11) of Sec. 6, Methods), predictions agree with the experimental measurements within errors (orange triangles). Results are reported in Extended Data Table 5. (**B**) Derivation of the entropic correction, $\delta\Delta s$. To do this, we subtracted the inverse of the measured, $T_m^{Bulk}$, and predicted, $T_m^{Unz}$, melting temperatures (blue squares). This equals the difference between the inverse of Eq.(11) and Eq.(10) (see Eq.(12) in Sec. 6, Methods). A linear fit to data (dashed red line) yields $\delta\Delta s = 6(1)\,\mathrm{cal\,mol^{-1}K^{-1}} \sim 4R\log 2$, where $R = 1.987\,\mathrm{cal\,mol^{-1}K^{-1}}$ is the ideal gas constant. The orange triangles show the theoretical correction to $T_m$ per DNA duplex predicted by assuming $\delta\Delta s \equiv 4R\log 2$.

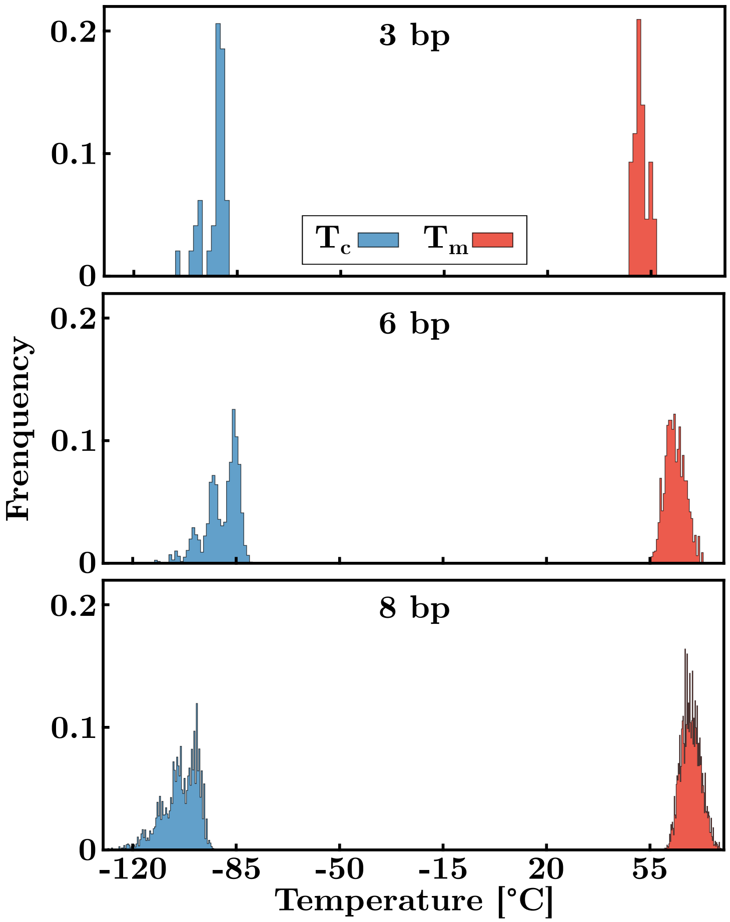


**Fig. 7**: **Prediction of DNA cold denaturation temperatures.** Histograms of the melting (red) cold denaturation (blue) temperatures predicted using the ten NNBP thermodynamics parameters (Table 1, main) for all possible DNA sequences of 3, 6, and 8 bp ending with a GAAA tetraloop.

### Extended Data: Tables

**Table 1: T-dependence of the DNA FDCs**

| T [°C] | T [K] | $f_m$ [pN] | $l_p$ [nm] | $d_b$ [nm] | $\Delta s_{ss}$ [cal mol$^{-1}$K$^{-1}$] |
|---|---|---|---|---|---|
| 7 | 280 | 19.72 (3) | 0.74 (7) | 0.647 (3) | 3.33 (2) |
| 10 | 283 | 19.08 (4) | 0.68 (2) | 0.631 (9) | 3.18 (2) |
| 13 | 286 | 18.71 (4) | 0.73 (3) | 0.662 (1) | 3.07 (2) |
| 16 | 289 | 18.26 (7) | 0.67 (2) | 0.672 (1) | 2.95 (2) |
| 19 | 292 | 17.87 (4) | 0.78 (2) | 0.655 (1) | 2.84 (2) |
| 22 | 295 | 17.12 (2) | 0.79 (3) | 0.657 (1) | 2.67 (2) |
| 25 | 298 | 16.75 (3) | 0.77 (2) | 0.647 (1) | 2.58 (2) |
| 30 | 303 | 15.86 (2) | 0.75 (2) | 0.665 (1) | 2.39 (2) |
| 35 | 308 | 14.96 (3) | 0.88 (2) | 0.639 (1) | 2.21 (1) |
| 42 | 315 | 14.06 (4) | 0.88 (4) | 0.641 (1) | 2.02 (1) |

FDC average unzipping force, $f_{\mathrm{m}}$, persistence length, $l_p$, interphosphate distance, $d_b$, and ssDNA stretching entropy per base, $\Delta s_{ss}$ in the studied temperature range (in Celsius and Kelvin degrees). The errors (in brackets) refer to the last digit. The error in temperature is ±1°C (K).

**Table 2**: NNBP $\Delta s_{0,i}$ $[\mathrm{cal\,mol^{-1}K^{-1}}]$ at different temperatures.

| **Temperature ±1 [K]** | **280** | **283** | **285** | **288** | **291** | **295** | **298** | **303** | **308** | **315** |
|---|---|---|---|---|---|---|---|---|---|---|
| **AA/TT** | -12.4 (5) | -12.6 (4) | -15.7 (7) | -12.8 (3) | -15.0 (5) | -16.0 (5) | -16.3 (6) | -16.9 (5) | -16.0 (3) | -18.3 (4) |
| **AC/TG** | -15.4 (2) | -15.6 (1) | -16.4 (2) | -16.4 (1) | -16.9 (2) | -17.5 (1) | -17.7 (1) | -18.3 (1) | -18.6 (1) | -19.6 (2) |
| **AG/TC** | -12.0 (3) | -13.1 (2) | -11.8 (4) | -14.3 (2) | -13.3 (4) | -13.8 (4) | -14.5 (3) | -15.0 (4) | -16.2 (3) | -16.6 (3) |
| **AT/TA** | -15.1 (1) | -15.3 (1) | -16.1 (1) | -15.6 (1) | -16.4 (1) | -17.1 (2) | -16.7 (1) | -17.3 (1) | -17.3 (1) | -17.9 (1) |
| **CA/GT** | -12.8 (2) | -13.7 (2) | -13.8 (2) | -14.9 (2) | -15.1 (2) | -16.6 (4) | -16.0 (2) | -17.3 (3) | -19.4 (7) | -18.4 (2) |
| **CC/GG** | -12.4 (3) | -13.3 (3) | -12.2 (4) | -14.6 (2) | -13.5 (4) | -13.7 (4) | -14.8 (3) | -15.2 (3) | -16.0 (3) | -16.4 (3) |
| **CG/GC** | -22.2 (3) | -22.4 (4) | -21.4 (5) | -23.8 (2) | -22.8 (4) | -22.6 (4) | -23.7 (3) | -24.1 (3) | -24.2 (4) | -24.6 (3) |
| **GA/CT** | -12.4 (4) | -13.4 (3) | -12.4 (5) | -14.6 (3) | -14.1 (4) | -14.5 (4) | -14.7 (5) | -15.2 (6) | -16.6 (4) | -17.3 (3) |
| **GC/CG** | -20.8 (3) | -21.3 (3) | -19.7 (5) | -22.8 (2) | -21.5 (4) | -21.8 (3) | -22.7 (3) | -23.5 (2) | -24.4 (3) | -23.7 (3) |
| **TA/AT** | -16.1 (1) | -16.1 (1) | -17.1 (2) | -16.2 (1) | -16.9 (1) | -17.2 (1) | -17.5 (2) | -17.7 (2) | -16.7 (3) | -18.2 (1) |

The 10 DNA entropies measured from unzipping a 3.6kbp hairpin in the temperature range $[280, 315]$ K (see text). The entropy of the last two motifs (GC/CG and TA/AT) has been computed by applying the circular symmetry relations. The error (in brackets) refers to the last digit.

**Table 3**: NNBP $\Delta g_{0,i}$ [kcal mol$^{-1}$] at different temperatures.

| **Temperature ±1 [K]** | **280** | **283** | **285** | **288** | **291** | **295** | **298** | **303** | **308** | **315** |
|---|---|---|---|---|---|---|---|---|---|---|
| **AA/TT** | −1.57(2) | −1.49(2) | −1.55(2) | −1.47(2) | −1.43(1) | −1.43(1) | −1.30(1) | −1.27(1) | −1.19(1) | −1.15(1) |
| **AC/TG** | −1.52(2) | −1.38(1) | −1.53(2) | −1.35(1) | −1.47(2) | −1.34(1) | −1.43(1) | −1.36(1) | −1.17(1) | −1.21(1) |
| **AG/TC** | −1.73(2) | −1.66(2) | −1.49(2) | −1.60(2) | −1.54(2) | −1.42(1) | −1.41(1) | −1.25(1) | −1.35(1) | −1.25(1) |
| **AT/TA** | −1.37(1) | −1.43(1) | −1.27(1) | −1.32(1) | −1.25(1) | −1.28(1) | −1.17(1) | −1.00(1) | −1.19(1) | −1.00(1) |
| **CA/GT** | −1.86(2) | −1.92(2) | −1.82(2) | −1.88(2) | −1.82(2) | −1.91(2) | −1.65(2) | −1.68(2) | −1.70(2) | −1.52(2) |
| **CC/GG** | −2.03(2) | −1.86(2) | −2.04(2) | −1.91(2) | −2.00(2) | −1.88(2) | −1.91(2) | −1.86(2) | −1.56(2) | −1.70(2) |
| **CG/GC** | −2.51(3) | −2.52(3) | −2.39(2) | −2.46(3) | −2.35(2) | −2.24(2) | −2.43(2) | −2.30(2) | −2.03(2) | −1.93(2) |
| **GA/CT** | −1.52(2) | −1.59(2) | −1.55(2) | −1.48(2) | −1.49(2) | −1.42(1) | −1.52(2) | −1.46(2) | −1.26(1) | −1.20(1) |
| **GC/CG** | −2.79(3) | −2.83(3) | −2.51(3) | −2.78(3) | −2.55(3) | −2.53(3) | −2.49(3) | −2.36(2) | −2.34(2) | −2.11(2) |
| **TA/AT** | −1.31(1) | −1.19(1) | −1.10(1) | −1.11(1) | −1.10(1) | −1.00(1) | −1.00(1) | −0.74(1) | −0.97(1) | −0.88(1) |

The 10 DNA free-energies measured from unzipping a 3.6kbp hairpin in the temperature range $[280, 315]$ K (see text). The entropy of the last two motifs (GC/CG and TA/AT) has been computed by the applying circular symmetry relations. The error (in brackets) refers to the last digit.

**Table 4**: NNBP $\Delta h_{0,i}$ [kcal mol$^{-1}$] at different temperatures.

| **Temperature ±1 [K]** | **280** | **283** | **285** | **288** | **291** | **295** | **298** | **303** | **308** | **315** |
|---|---|---|---|---|---|---|---|---|---|---|
| **AA/TT** | -5.05 (15) | -5.04 (12) | -6.04 (21) | -5.16 (08) | -5.83 (16) | -6.17 (15) | -6.16 (17) | -6.39 (15) | -6.13 (09) | -6.93 (14) |
| **AC/TG** | -5.84 (05) | -5.80 (03) | -6.21 (05) | -6.10 (04) | -6.41 (05) | -6.51 (04) | -6.71 (05) | -6.91 (04) | -6.90 (03) | -7.38 (06) |
| **AG/TC** | -5.09 (10) | -5.36 (07) | -4.86 (13) | -5.74 (06) | -5.41 (12) | -5.51 (12) | -5.73 (10) | -5.80 (12) | -6.34 (09) | -6.49 (09) |
| **AT/TA** | -5.60 (03) | -5.76 (03) | -5.88 (04) | -5.82 (03) | -6.03 (04) | -6.32 (05) | -6.15 (03) | -6.26 (04) | -6.54 (04) | -6.65 (03) |
| **CA/GT** | -5.45 (05) | -5.81 (05) | -5.76 (06) | -6.19 (06) | -6.22 (07) | -6.81 (11) | -6.41 (05) | -6.93 (09) | -7.67 (20) | -7.31 (06) |
| **CC/GG** | -5.51 (10) | -5.62 (08) | -5.52 (12) | -6.13 (06) | -5.93 (11) | -5.93 (12) | -6.32 (08) | -6.46 (10) | -6.51 (10) | -6.88 (08) |
| **CG/GC** | -8.75 (09) | -8.86 (11) | -8.49 (14) | -9.32 (07) | -9.01 (11) | -8.92 (13) | -9.51 (09) | -9.60 (09) | -9.50 (12) | -9.68 (10) |
| **GA/CT** | -4.99 (11) | -5.37 (08) | -5.09 (14) | -5.71 (08) | -5.59 (11) | -5.70 (13) | -5.91 (14) | -6.07 (17) | -6.38 (12) | -6.63 (11) |
| **GC/CG** | -8.61 (10) | -8.85 (10) | -8.15 (14) | -9.38 (06) | -8.82 (11) | -8.98 (11) | -9.27 (09) | -9.48 (07) | -9.87 (09) | -9.57 (11) |
| **TA/AT** | -5.84 (04) | -5.75 (03) | -5.99 (05) | -5.79 (03) | -6.04 (04) | -6.07 (04) | -6.21 (05) | -6.11 (05) | -6.14 (08) | -6.61 (05) |

The 10 DNA enthalpies measured from unzipping a 3.6kbp hairpin in the temperature range $[280, 315]$ K (see text). The entropy of the last two motifs (GC/CG and TA/AT) has been computed by the applying circular symmetry relations. The error (in brackets) refers to the last digit.

**Table 5**: DNA Oligos Melting Temperatures [°C]

| **Sequence** ($5' \rightarrow 3'$) | $T^{Exp}$ | $T^{Unz}_{Bi}$ | $T^{Unz}_{Uni}$ | $T^{UO}$ | $T^{Hug}$ |
|---|---|---|---|---|---|
| ATCAATCATA | 33.6 | 40.6 | 33.1 | 34.0 | 32.3 |
| TTGTAGTCAT | 36.0 | 42.4 | 35.0 | 36.7 | 36.2 |
| GAAATGAAAG | 34.4 | 44.6 | 37.3 | 34.6 | 33.8 |
| CCAACTTCTT | 40.6 | 47.9 | 40.6 | 40.4 | 38.9 |
| ATCGTCTGGA | 44.9 | 49.6 | 42.1 | 46.2 | 45.3 |
| AGCGTAAGTC | 40.3 | 48.2 | 40.9 | 45.1 | 43.4 |
| CGATCTGCGA | 49.1 | 54.5 | 47.3 | 50.5 | 50.2 |
| TGGCGAGCAC | 55.3 | 59.5 | 52.4 | 56.3 | 54.7 |
| GATGCGCTCG | 53.5 | 57.7 | 50.6 | 54.0 | 52.9 |
| GGGACCGCCT | 57.0 | 59.8 | 52.3 | 58.6 | 55.3 |
| CGTACACATGC | 49.9 | 53.8 | 47.3 | 51.2 | 50.6 |
| CCATTGCTACC | 48.9 | 55.5 | 48.7 | 49.6 | 48.2 |
| TACTAACATTAACTA | 51.1 | 54.6 | 49.3 | 51.9 | 50.1 |
| ATACTTACTGATTAG | 51.5 | 56.0 | 50.6 | 49.7 | 50.4 |
| GTACACTGTCTTATA | 54.8 | 56.7 | 51.4 | 54.8 | 54.2 |
| GTATGAGAGACTTTA | 55.4 | 58.5 | 53.2 | 54.8 | 54.2 |
| TTCTACCTATGTGAT | 53.7 | 60.3 | 55.0 | 55.1 | 55.7 |
| AGTAGTAATCACACC | 57.1 | 59.8 | 54.5 | 56.9 | 57.2 |
| ATCGTCTCGGTATAA | 58.6 | 61.8 | 56.5 | 58.9 | 59.9 |
| ACGACAGGTTTACCA | 61.3 | 65.3 | 60.1 | 63.6 | 63.6 |
| CTTTCATGTCCGCAT | 62.8 | 68.0 | 62.9 | 63.0 | 62.6 |
| TGGATGTGTGAACAC | 60.4 | 64.8 | 59.8 | 62.3 | 62.1 |
| ACCCCGCAATACATG | 62.9 | 68.9 | 63.8 | 65.6 | 64.5 |
| GCAGTGGATGTGAGA | 63.3 | 68.1 | 63.0 | 64.6 | 64.2 |
| GGTCCTTACTTGGTG | 60.3 | 65.2 | 60.0 | 62.0 | 61.7 |
| CGCCTCATGCTCATC | 65.8 | 70.9 | 65.8 | 66.5 | 65.9 |
| AAATAGCCGGGCCGC | 70.4 | 75.8 | 70.7 | 72.7 | 70.9 |
| CCAGCCAGTCTCTCC | 66.7 | 70.9 | 65.7 | 67.7 | 66.7 |
| GACGACAAGACCGCG | 68.6 | 69.7 | 64.7 | 69.7 | 70.3 |
| CAGCCTCGTCGCAGC | 72.0 | 74.8 | 69.8 | 73.0 | 72.7 |
| CTCGCGGTCGAAGCG | 70.7 | 73.7 | 68.7 | 72.9 | 73.6 |
| GCGTCGGTCCGGGCT | 74.1 | 76.5 | 71.4 | 77.8 | 76.1 |
| TATGTATATTTTGTAATCAG | 61.2 | 64.9 | 60.8 | 58.6 | 59.5 |
| TTCAAGTTAAACATTCTATC | 61.5 | 67.6 | 63.6 | 60.6 | 62.6 |
| TGATTCTACCTATGTGATTT | 64.4 | 69.5 | 65.4 | 63.7 | 64.8 |
| GAGATTGTTTCCCTTTCAAA | 65.3 | 72.8 | 68.8 | 66.3 | 67.1 |
| ATGCAATGCTACATATTCGC | 68.9 | 74.7 | 70.8 | 69.2 | 69.6 |
| CCACTATACCATCTATGTAC | 64.4 | 67.5 | 63.4 | 63.9 | 65.2 |
| CCATCATTGTGTCTACCTCA | 68.5 | 73.0 | 69.0 | 69.4 | 69.7 |
| CGGGACCAACTAAAGGAAAT | 68.5 | 74.5 | 70.5 | 70.3 | 70.5 |
| TAGTGGCGATTAGATTCTGC | 71.2 | 74.6 | 70.6 | 71.1 | 70.9 |
| AGCTGCAGTGGATGTGAGAA | 73.1 | 78.0 | 74.1 | 74.5 | 74.0 |
| TACTTCCAGTGCTCAGCGTA | 73.6 | 76.2 | 72.2 | 76.0 | 74.6 |
| CAGTGAGACAGCAATGGTCG | 72.5 | 76.0 | 72.1 | 73.5 | 73.9 |
| CGAGCTTATCCCTATCCCTC | 70.3 | 75.5 | 71.4 | 71.3 | 71.9 |
| CGTACTAGCGTTGGTCATGG | 71.1 | 74.6 | 70.7 | 72.9 | 74.0 |
| AAGGCGAGTCAGGCTCAGTG | 76.3 | 79.4 | 75.5 | 77.2 | 77.0 |
| ACCGACGACGCTGATCCGAT | 77.3 | 78.3 | 74.4 | 78.7 | 78.6 |
| AGCAGTCCGCCACACCCTGA | 78.5 | 82.0 | 78.1 | 81.6 | 79.7 |
| CAGCCTCGTTCGCACAGCCC | 78.1 | 82.1 | 78.3 | 81.1 | 80.4 |
| GTGGTGGGCCGTGCGCTCTG | 81.0 | 83.2 | 79.4 | 83.6 | 82.0 |
| GTCCACGCCCGGTGCGACGG | 81.1 | 83.0 | 79.1 | 85.4 | 84.2 |
| GATATAGCAAAATTCTAAGTTAATA | 66.1 | 71.5 | 68.2 | 64.2 | 65.9 |
| ATAACTTTACGTGTGTGACCTATTA | 71.8 | 73.5 | 70.2 | 71.2 | 72.4 |
| GTTCTATACTCTTGAAGTTGATTAC | 67.7 | 72.2 | 68.9 | 67.3 | 69.9 |
| CCCTGCACTTTAACTGAATTGTTTA | 72.5 | 77.7 | 74.5 | 73.4 | 73.5 |
| TAACCATACTGAATACCTTTTGACG | 71.3 | 75.4 | 72.1 | 72.2 | 73.0 |
| TCCACACGGTAGTAAAATTAGGCTT | 73.8 | 78.2 | 74.9 | 74.6 | 75.8 |
| TTCCAAAAGGAGTTATGAGTTGCGA | 73.8 | 79.8 | 76.6 | 75.2 | 76.2 |
| AATATCTCTCATGCGCCAAGCTACA | 76.5 | 81.3 | 78.1 | 76.7 | 77.3 |
| TAGTATATCGCAGCATCATACAGGC | 75.0 | 78.8 | 75.5 | 75.5 | 75.8 |
| TGGATTCTACTCAACCTTAGTCTGG | 73.6 | 77.8 | 74.5 | 73.9 | 75.2 |
| CGGAATCCATGTTACTTCGGCTATC | 74.8 | 79.0 | 75.7 | 75.5 | 76.6 |

**Table 5**: DNA Oligos Melting Temperatures [°C]

| **Sequence** ($5' \rightarrow 3'$) | $\mathbf{T^{Exp}}$ | $\mathbf{T^{Unz}_{Bi}}$ | $\mathbf{T^{Unz}_{Uni}}$ | $\mathbf{T^{UO}}$ | $\mathbf{T^{Hug}}$ |
|---|---|---|---|---|---|
| CTGGTCTGGATCTGAGAACTTCAGG | 75.6 | 80.1 | 76.8 | 76.6 | 77.3 |
| ACAGCGAATGGACCTACGTGGCCTT | 81.0 | 83.6 | 80.4 | 82.7 | 82.1 |
| AGCAAGTCGAGCAGGGCCTACGTTT | 81.5 | 84.5 | 81.3 | 82.8 | 82.8 |
| GCGAGCGACAGGTTACTTGGCTGAT | 80.1 | 83.1 | 79.9 | 81.3 | 81.7 |
| AAAGGTGTCGCGGAGAGTCGTGCTG | 82.4 | 83.5 | 80.4 | 83.0 | 83.6 |
| ATGGGTGGGAGCCTCGGTAGCAGCC | 83.4 | 87.4 | 84.1 | 86.6 | 84.6 |
| CAGTGGGCTCCTGGGCGTGCTGGTC | 83.4 | 87.6 | 84.4 | 87.5 | 85.7 |
| GCCAACTCCGTCGCCGTTCGTGCGC | 84.6 | 86.9 | 83.8 | 88.1 | 88.0 |
| ACGGGTCCCCGCACCGCACCGCCAG | 88.3 | 90.4 | 87.2 | 93.0 | 90.1 |
| TTATGTATTAAGTTATATAGTAGTAGTAGT | 66.6 | 71.4 | 68.5 | 65.8 | 69.7 |
| ATTGATATCCTTTTCTATTCATCTTTCATT | 70.4 | 78.0 | 75.2 | 70.3 | 71.8 |
| AAAGTACATCAACATAGAGAATTGCATTTC | 73.2 | 78.8 | 76.1 | 73.0 | 74.6 |
| CTTAAGATATGAGAACTTCAACTAATGTGT | 71.8 | 77.1 | 74.3 | 71.8 | 74.2 |
| CTCAACTTGCGGTAAATAAATCGCTTAATC | 75.5 | 80.5 | 77.8 | 75.2 | 77.3 |
| TATTGAGAACAAGTGTCCGATTAGCAGAAA | 76.4 | 81.2 | 78.5 | 77.5 | 78.4 |
| GTCATACGACTGAGTGCAACATTGTTCAAA | 76.9 | 80.8 | 78.2 | 78.0 | 79.3 |
| AACCTGCAACATGGAGTTTTTGTCTCATGC | 78.7 | 83.7 | 81.1 | 80.3 | 80.1 |
| CCGTGCGGTGTGTACGTTTTATTCATCATA | 77.6 | 81.2 | 78.5 | 80.0 | 80.5 |
| GTTCACGTCCGAAAGCTCGAAAAAGGATAC | 78.7 | 82.1 | 79.4 | 79.5 | 81.5 |
| AGTCTGGTCTGGATCTGAGAACTTCAGGCT | 80.6 | 84.7 | 81.9 | 82.2 | 82.5 |
| TCGGAGAAATCACTGAGCTGCCTGAGAAGA | 80.9 | 86.0 | 83.3 | 82.5 | 83.3 |
| CTTCAACGGATCAGGTAGGACTGTGGTGGG | 80.1 | 84.4 | 81.7 | 83.3 | 83.4 |
| ACGCCCACAGGATTAGGCTGGCCCACATTG | 84.0 | 88.9 | 86.2 | 87.5 | 85.5 |
| GTTATTCCGCAGTCCGATGGCAGCAGGCTC | 84.1 | 87.8 | 85.1 | 85.9 | 85.6 |
| TCAGTAGGCGTGACGCAGAGCTGGCGATGG | 84.6 | 88.8 | 86.1 | 88.1 | 88.2 |
| CGCGCCACGTGTGATCTACAGCCGTTCGGC | 84.5 | 88.2 | 85.6 | 89.0 | 89.3 |
| GACCTGACGTGGACCGCTCCTGGGCGTGGT | 86.4 | 89.3 | 86.6 | 91.2 | 89.9 |
| GCCCCTCCACTGGCCGACGGCAGCAGGCTC | 87.7 | 93.3 | 90.6 | 93.8 | 91.5 |
| CGCCGCTGCCGACTGGAGGAGCGCGGGACG | 88.6 | 93.4 | 90.8 | 94.8 | 93.9 |

Melting temperatures of the 92 DNA duplexes studied by Owczarzy *et al.* in Ref. [55] at a concentration $c = 2\mu$M and 1020mM NaCl. The experimental values ($T^{\rm Exp}$) are compared with predictions obtained with the unzipping parameters by using Eq.(10) ($T^{\rm Unz}_{\rm Bi}$) and Eq.(11) ($T^{\rm Unz}_{\rm Uni}$) for bimolecular and unimolecular reactions, respectively (see main text and Sec. 6, Methods). Finally, $T^{\rm UO}$ and $T^{\rm Hug}$ are obtained with the unified oligonucleotide parameters (UO) in Ref. [43] and the Huguet *et al.* (2017) parameters in Ref. [37]. Results are reported with errors: $T^{\rm Exp} \pm 1.6$°C, $T^{\rm Unz} \pm 1.5$°C, $T^{\rm UO} \pm 1.5$°C, and $T^{\rm Hug} \pm 1.5$°C. Temperatures are given in Celsius degrees.
